# A protein phosphorylation module patterns the *Bacillus subtilis* spore outer coat

**DOI:** 10.1101/469122

**Authors:** Carolina Freitas, Jerneja Plannic, Rachele Isticato, Assunta Pelosi, Rita Zilhão, Mónica Serrano, Loredana Baccigalupi, Ezio Ricca, Alexander K.W. Elsholz, Richard Losick, Adriano O. Henriques

## Abstract

Assembly of the *Bacillus subtilis* spore coat involves over 80 protein components, which self-organize into a basal layer, a lamellar inner coat, a striated electrondense outer coat and a more external crust. CotB is an abundant component of the outer coat. Its C-terminal moiety contains a region, termed SKR^B^, formed by a series of serine-rich repeats, which we show is phosphorylated by the coat-associated Ser/Thr kinase CotH at multiple Ser residues. Another coat protein, CotG, which contains a central repeat region, SKR^G^, interacts with the C-terminal moiety of CotB and promotes its phosphorylation by CotH *in vivo* and in a heterologous system. CotG itself is phosphorylated by CotH but phosphorylation is enhanced in the absence of CotB. Spores of a *cotH*^*D288A*^ strain, producing an inactive form of the kinase, like those formed by a *cotG* deletion mutant, lack the characteristic pattern of electrondense outer coat striations, while retaining the crust. Specifically, in the absence of CotG or CotH activity, most of the outer coat proteins are assembled but form a layer of amorphous material that peels-off the spore if crust formation is genetically ablated. In contrast, deletion of the SKR^B^ region, has no major impact on the structure of the outer coat. Thus, phosphorylation of CotG by CotH is the principal factor establishing the structural pattern of the spore outer coat. The presence of the *cotB*/*cotH*/*cotG* cluster in several species closely related to *B. subtilis* and of *cotG*-like proteins in nearly all spore-formers that also code for a CotH homologue hints at the importance of this protein phosphorylation module in the morphogenesis of the spore outer layers.

## Introduction

Bacterial endospores (spores for simplicity) are a dormant cell type formed by a diverse group of bacteria within the Firmicutes phylum. Sporulation occurs within a sporangium formed by a larger mother cell, and a smaller forespore, or future spore. At the end of the differentiation process, and upon lysis of the mother cell, the spore is released into the environment. In *Bacillus subtilis*, the outermost spore layer is the coat, a protein-bound organelle that protects mature spores and mediates their interaction with abiotic and biotic surfaces and also with germinants (Driks & Eichenberger, 2016, Henriques & Moran, 2007, McKenney *et al.*, 2013). The coat comprises a basal layer, a lamellar inner coat, a striated elecontrondense outer coat and a more external crust (Driks & Eichenberger, 2016, Henriques & Moran, 2007, McKenney *et al.*, 2013). In species of the *B. cereus*/*B. anthracis*/*B. thuringiensis* group, the more external spore layer is an exosporium, formed by a basal layer and a hair-like nap that projects from it; this layer is separated from the coat by an interspace of variable length (Stewart, 2015).

Synthesis of the coat and crust proteins relies on a mother cell type-specific transcriptional cascade involving two RNA polymerase sigma factors and three ancillary transcription factors, in the order σ^E^, SpoIIID and GerR, σ^K^, and GerE (Driks & Eichenberger, 2016, Henriques & Moran, 2007, McKenney *et al.*, 2013). The morphogenetic proteins that govern basal layer (SpoIVA and SpoVM), inner coat (SafA), outer coat (CotE) and crust assembly (CotZ), are produced early in development, when the larger mother cell begins engulfment of the smaller forespore, and are recruited to the mother cell proximal forespore pole (MCP) to form an organizational center responsible for the assembly of the various coat sub-layers (McKenney *et al.*, 2013). In a second step in coat assembly, termed encasement, the coat proteins start surrounding the forespore, some tracking the engulfing membranes, others in successive waves during and following engulfment completion; the waves in encasement are determined by the deployment of the mother cell transcriptional cascade (McKenney *et al.*, 2013). Self-assembly mechanisms and post-translational modifications of the coat proteins such as proteolytical processing, glycosylation or cross-linking that also play important roles in coat assembly and maturation (Driks & Eichenberger, 2016, McKenney *et al.*, 2013). SpoIVA, for example, is an ATPase that self-assembles into cables, in an ATP-dependent manner, that cover the surface of the forespore to form the coat basal layer (Ramamurthi & Losick, 2008). An important question is how the structural features of the spore surface layers arise from individual components and recent work has shed some light onto the assembly of the crust in *B. subtilis* and the exosporium in *B. cereus*/*B. anthracis.* Crust and exosporium proteins self assemble, in this case to form into two-dimensional sheets with a hexagonal lattice and three-dimensional stacks which reproduce structures seen in mature spores (Ball *et al.*, 2008, Jiang *et al.*, 2015, Kailas *et al.*, 2011). However, how the structural features of the coat layers emerge is unknown.

Here, we are concerned with the formation of the electrondense striated outer coat layer of *B. subtilis* spores. CotB and CotG are two abundant outer coat components (Donovan *et al.*, 1987, Sacco *et al.*, 1995, Zilhao *et al.*, 2004) and an earlier study has suggested that CotG is an important structural determinant of this layer (Henriques *et al.*, 1998). The *cotB* and *cotG* genes are clustered together with a third gene, *cotH*, which is also an important determinant of outer coat assembly (Kim *et al.*, 2006, Naclerio *et al.*, 1996, Sacco *et al.*, 1995, Saggese *et al.*, 2014, Zilhao *et al.*, 1999) (Fig. 1A) and codes for a eukaryotic-type Ser/Thr kinase (Galperin *et al.*, 2012, Nguyen *et al.*, 2016). Both CotB and CotG possess a series of continuous Ser/Lys/Arg-rich tandem repeats, CotB in its C-terminal moiety (the SKR^B^ region) and CotG in its central part (the SKR^G^ region) (Sacco *et al.*, 1995, Saggese *et al.*, 2014) (Fig. 1B and C). Although CotB is synthesized as a 46 kDa species (CotB-46) the main form of the protein detected in mature spores has an apparent mass of 66 kDa (CotB-66) (Naclerio *et al.*, 1996, Sacco *et al.*, 1995, Zilhao *et al.*, 2004). Accumulation of CotB-66 in spores requires both CotH and CotG (Zilhao *et al.*, 2004). It is likely that CotB-66 is a polyphosphorylated form of the protein; not only is CotB-46 phosphorylated by CotH in vitro to produce a species with an apparent mass of about 66 kDa, but a phosphorylated protein at about 66 kDa is detected in extracts from WT spores but not in those obtained from a *cotH* mutant (Nguyen *et al.*, 2016). CotG is also phosphorylated in the SKR^G^ region during spore coat assembly and a peptide derived from SKR^G^ is phosphorylated by CotH in vitro (Nguyen *et al.*, 2016, Saggese *et al.*, 2014). Thus, the kinase activity of CotH is required for the phosphorylation of CotG and CotB, and possibly of additional spore coat proteins (Nguyen *et al.*, 2016).

**Figure 1.**
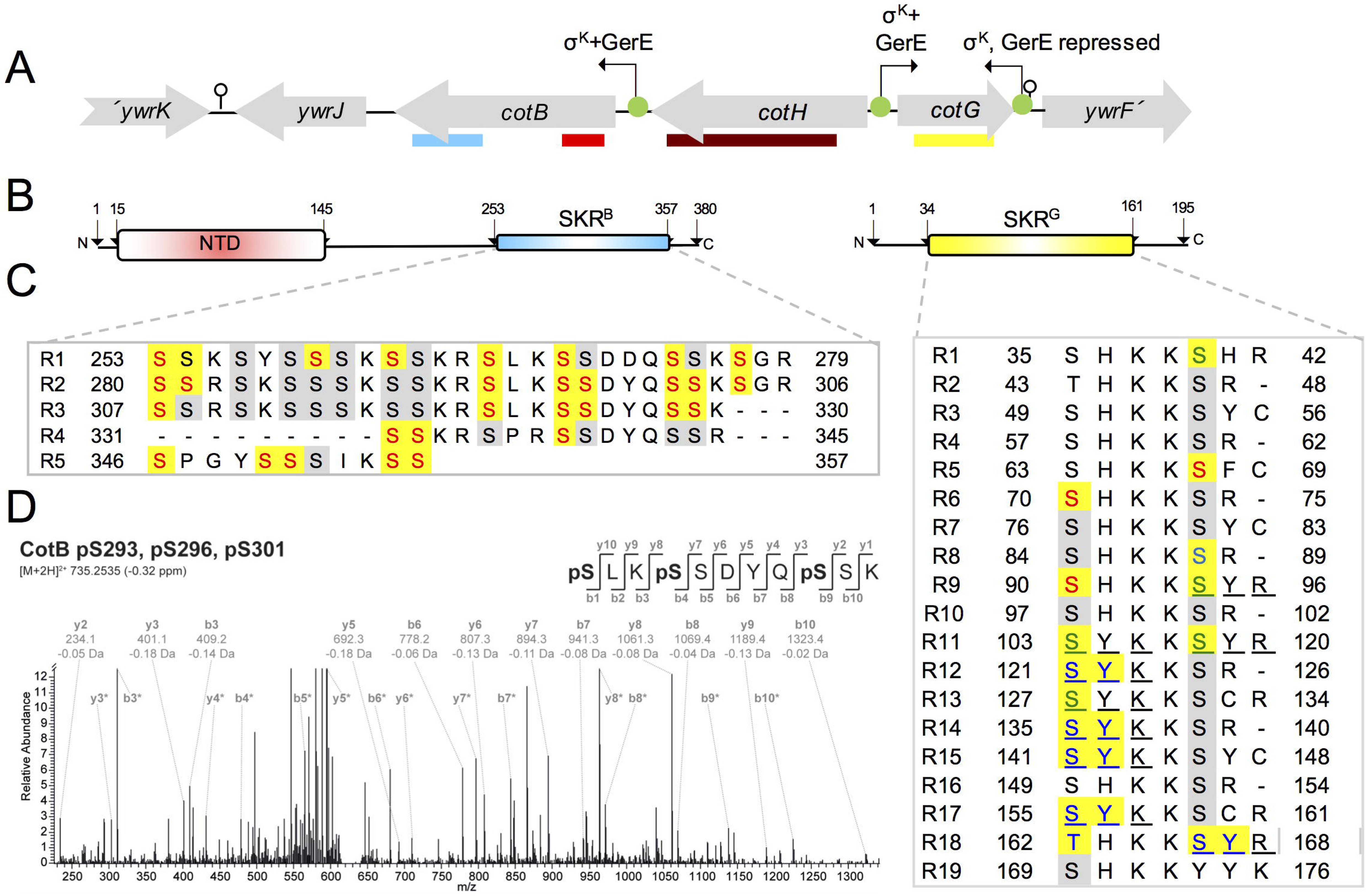
The *cotBGH* cluster and phosphosites in CotB and CotG. **A:** the *cotB, cotG* and *cotH* region of the *B. subtilis* chromosome. The red line below *cotB* delimits a N-terminal domain (NTD) and the blue line represents the SKR^B^ region. The yellow line below the *cotG* gene shows the position of the SKR^G^ region (Giglio *et al.*, 2011). The brown line below *cotH* shows the region of homology to eukaryotic-type Ser/Thr kinases (Nguyen *et al.*, 2016). Promoters are represented by the green circles and broken arrows, and their known regulators are indicated. The stem and loop structures represent putative transcriptional terminators. **B**: structural organization of CotB (left) and CotG (right), with the NTD in CotB and the SKR^B^ and SKR^G^ regions in evidence. **C**: Phosphorylated residues in CotB and CotG. The sequence of the repeats in the SKR^B^ and SKR^G^ regions is shown, with conserved Ser residues highlighted against a grey or yellow (when phosphorylated) background. pSer residues found for CotB in our study are in red; the pSer residue detected in the proteome of germinating spores is shown in black against the yellow background (Rosenberg *et al.*, 2015). pSer residues in CotG are indicated as follows: those detected in our study are in red, those found in the study of Saggese *et al.* (Saggese *et al.*, 2014) are in blue and those detected in both studies are in green. Tripeptides containing a phosphate moiety found by Saggese *et al*. (Saggese *et al.*, 2014) are underlined. In some cases, pSer was unambiguously identified in those same tripeptides in our study (green). Ser88 or Ser90, in the context of a peptide (residues 88-100) were phosphorylated by purified CotH (Nguyen *et al.*, 2016). **D**: Identification of multiple serine phosphorylation sites in CotB with HPLC-MS/MS analyses of phosphopeptide-enriched samples. The fragment spectra generated by collision-induced dissociation of the tryptic phosphopeptide precursor pSLKpSSDYQpSSK at m/z 735.3 (M+2H)2+ leading to the identification of phosphoserine at positions 293, 296, and 301, within repeat R2, is shown. Signals assigned to fragments of the b-and y-ion series are labelled with the theoretical mass and the delta mass between theoretical and observed mass. Additional b and y-ion signals resulting from neutral loss of phosphoric acid (-H_3_PO_4_) are indicated with asterisk.

The *cotB*/*cotH*/*cotG* cluster is found in several species closely related to *B. subtilis* and also in *Geobacillus* (McPherson *et al.*, 2010, Naclerio *et al.*, 1996, Sacco *et al.*, 1995, Saggese *et al.*, 2016, Todd *et al.*, 2003). While CotB homologues are found in most spore-forming Bacilli, non-homologous CotG-like proteins are found in nearly all spore-forming species that also code for a CotH homologue (Galperin *et al.*, 2012, Saggese *et al.*, 2016). In the *B. cereus* group, for instance, ExsB, a CotG-like protein required for attachment of the exosporium to the coat, is phosphorylated at multiple Thr residues within a central repeat region; this phosphorylation most likely relies on a CotH homologue (McPherson *et al.*, 2010, Nguyen *et al.*, 2016). Importantly, CotG was also proposed to be a crust component in *B. subtilis* (McPherson *et al.*, 2010). Two CotB paralogues are also found in the coat/exosporia of *B. cereus*/*B. anthracis* spores (Abhyankar *et al.*, 2013, Abhyankar *et al.*, 2017), suggesting that the *cotB*/*cotG*/*cotH* cluster may participate in determining the the structural pattern seen in spores of these organisms: a thin outer coat/interspace/exosporium versus the thick outer coat/crust architecture of *B. subtilis* spores.

Previous studies on the structural characterization of the *cotB/cotH/cotG* cluster in *B. subtilis* made use of polar insertional mutations, and it is unclear how the absence of active CotH impacts on the overall structure of the coat and what is the contribution of phosphorylated CotG and CotB. Also unclear is how CotG influences the phosphorylation of CotB-46. We now show that CotG is required for the efficient phosphorylation of CotB-46 by CotH, both during coat assembly and also in a heterologous system. Conversely, phosphorylation of CotG in a CotH-dependent manner is favored in the absence of CotB-46. We show that CotG interacts with the C-terminal region of CotB and we propose that this interaction promotes phosphorylation of CotB-46 while inhibiting the phosphorylation of CotG in the presence of CotH. We show that phosphorylation of CotB-46 and CotG occurs at the surface of the developing spore. While deletion of the SKR^B^ region has no impact on the structure of the coat, we show that mutants lacking active CotH or CotG form an amorphous outer coat which is still delimited at its outer edge by the crust. Thus, CotH and CotG establish the normal striated pattern of the outer coat. Strikingly, the outer coat region of *cotH* or *cotG* mutants resemble the interspace and exosporium of *B. cereus*/*B. anthracis* spores suggesting that CotH and CotG may have a role in determining the structural and functional features of the spore surface layers in a wide range of organisms.

## Results

### Phosphorylation sites in CotB and CotG

A striking feature of CotB and CotG is the presence in the C-terminal half of CotB and in the central part of CotG, of direct repeats of a sequence rich in Ser, Lys, and Arg residues (Giglio *et al.*, 2011, Sacco *et al.*, 1995, Saggese *et al.*, 2016) (Fig. 1 and S1). The SKR^B^ region, in CotB, is formed by 4 direct repeats whereas SKR^G^, in CotG, is formed by 19 direct repeats of 5 to 7 amino acids (Giglio *et al.*, 2011, Sacco *et al.*, 1995, Saggese *et al.*, 2016). Both SKR^B^ and SKR^G^ are likely to be disordered (Fig. S1) (Liu & Huang, 2014, Romero *et al.*, 1997). Moreover, the Ser residues within SKR^B^ and SKR^G^, but much less so outside this region, show a high probability of being targets for phosphorylation (Fig. S1B) and indeed, several phosphorylation sites have been identified in both regions ((Giglio *et al.*, 2011, Nguyen *et al.*, 2016); see also below) (Fig. 1C; see also the supplemental text).

In our study, both CotB and CotG were detected as phosphorylated species by mass spectrometry of lysates prepared from sporulating cells. After induction of sporulation we collected samples every hour for five hours and proteins were digested using trypsin; the resulting peptides were enriched for phosphopeptides using TiO_2_ chromatography, followed by subsequent analysis using nano-liquid chromatography-tandem mass spectrometry (nanoLC-MS/MS) on Orbitrap mass spectrometers (see methods). pSer was identified at 29 positions in CotB, all of which are located within the SKR^B^ region (Fig. 1C). Phosphorylation of Ser253 and Ser254, the later not detected in our study, was also detected in CotB when purified spores were induced to germinate (Rosenberg *et al.*, 2015). Thus, CotB is phosphorylated at a minimum of 29-30 Ser residues within the SKR^B^ region (Fig. 1C). To date, site-specific phosphorylation of CotB was only shown for a single phosphosite per molecule, leaving the possibility that CotB can be phosphorylated at several serine residues, but only once per molecule. However, the dramatic shift of CotB-46 to CotB-66 in the presence of CotG and CotH suggests that CotB is polyphosphorlyated. Here, we were able to show multiple serine phosphosites in a single peptide, demonstrating the occurrence of multiple phosphosites within one CotB molecule (Fig. 1D; see also Table S4).

### The kinase activity of CotH explains its morphogenetic role

Using a phosphospecific antibody raised against the consensus phosphorylation site recognized by protein kinase C (PKC), Nguyen and co-authors showed the presence of phosphorylated proteins in spore coat extracts (Nguyen *et al.*, 2016). Bands of about 30 kDa, inferred to be CotG, and of 66 kDa inferred to correspond to CotB-66, were detected in extracts from WT spores, and in spores formed by a *cotH* mutant expressing WT *cotH in trans*; these bands were not detected in spores producing catalytic inactive forms of CotH from the same ectopic site in the chromosome, or in spores of a *cotH* null mutant (Nguyen *et al.*, 2016). Additional bands were detected, suggesting that CotH could phosphorylate other coat proteins (Nguyen *et al.*, 2016). CotH is known to control the assembly of at least nine major coat proteins (Giorno *et al.*, 2007, Naclerio *et al.*, 1996, Zilhao *et al.*, 1999). However, in the study of Nguyen and co-workers, a Coomassie-stained gel of coat extracts was not shown, precluding an evaluation of the impact of the catalytic mutants on the overall assembly of the coat. To more directly test the impact of CotH activity on coat assembly, we first constructed a strain producing a catalytic inactive form of CotH. Previous work has shown that the substitution of Asp228 in CotH by a Gln, renders the protein inactive ((Nguyen *et al.*, 2016); see also below). Here, the WT or a *cotH*^*D228Q*^ alleles were inserted at the non-essential *amyE* locus in a *cotH* insertional mutant; both alleles were placed under the control of the σ^K^-dependent promoter known to drive expression of *cotH* and which is located downstream of the 3′-end of *cotG* (Giglio *et al.*, 2011) (Fig. 1A and 2A). To avoid insertion of a second copy of *cotG* at *amyE*, the region corresponding to the long 5′-unstranslated region of the *cotH* mRNA was not included in our construct (Fig. 2A). Deletion of this region, however, does not affect the accumulation of CotH or spore coat formation (Giglio *et al.*, 2011).

**Figure 2.**
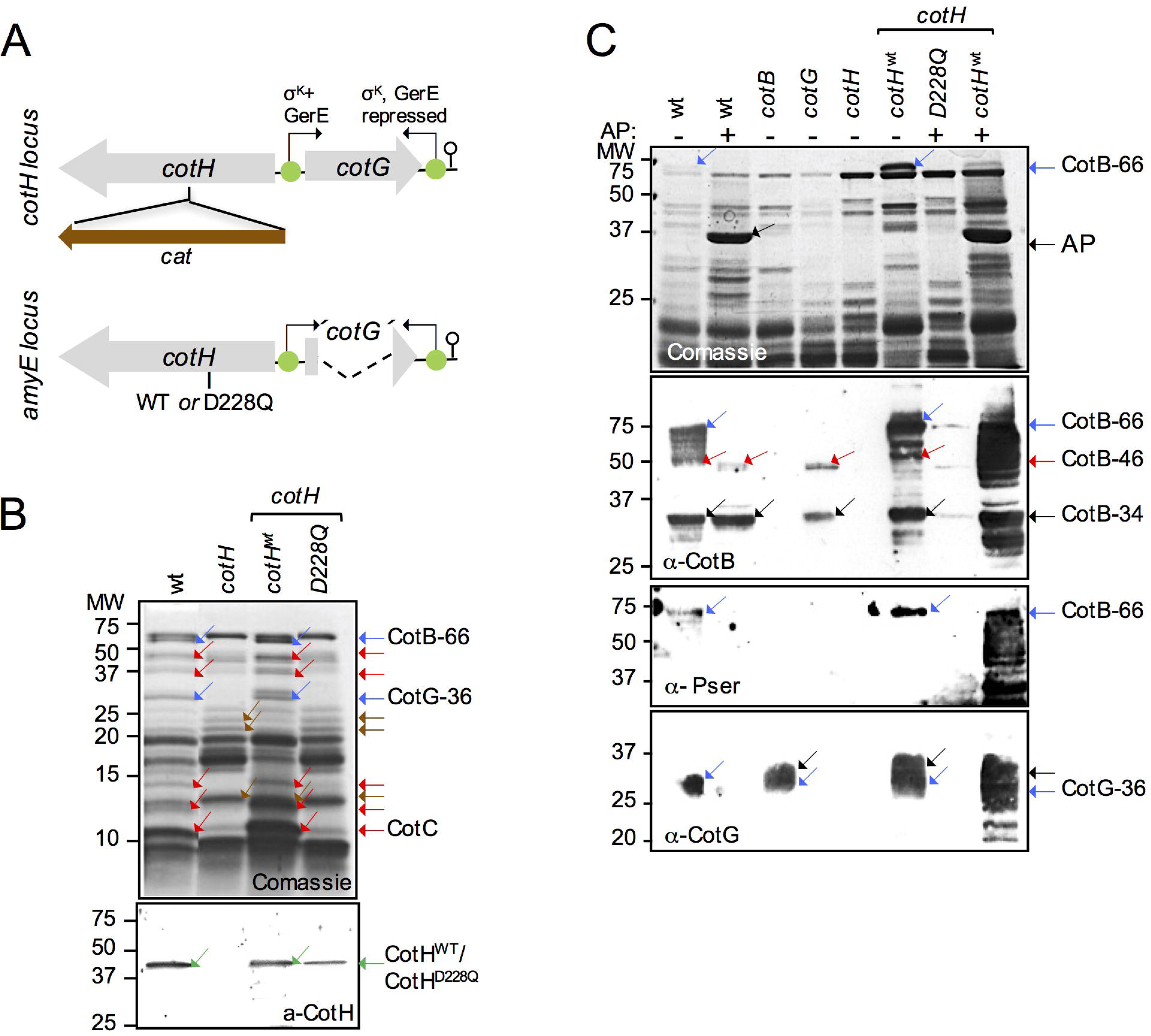
Phosphorylation of CotB-46 in spores requires CotG. **A:** construction of a mutant producing the catalytically inactive variant CotH^D228Q^. A chloramphenicol-resistance (*cat*) cassette was inserted in the *cotH* locus (top) and the WT or *D228Q* allelles of *cotH* were inserted into the *amyE* locus of the resulting strain. Both alleles carry an in-frame deletion of the *cotG* coding region and are under the control of the native *cotH* promoter (located downstream of the 3′-end of the *cotG* gene; green circles and broken arrow). **B**: shows spore coat proteins extracted from purified spores of the WT, Δ*cotH*, Δ*cotH*/*cotH*^*wt*^ and Δ*cotH*/*cotH*^*D228Q*^ strains, as indicated, using a NaOH treatment. The extracted proteins were resolved by SDS-PAGE (on 15% gels). The gel was stained with Coomassie (top panel) and subject to immunoblotting with an anti-CotH antibody (bottom). Red arrows, proteins showing reduced extractability in the *cotH* mutants; brown arrows; proteins with increased extractability in the mutants. **C:** Spores produced by a wild type strain, and *cotB, cotG*, and *cotH* mutants were purified, the coat proteins extracted by NaOH treatment, analysed by SDS-PAGE and Coomassie staining, and subject to immunobloting with anti-CotB, anti-phospho-Serine or anti-CotG antibodies, as indicated. Proteins extracted from wild type spores were analysed before and after treatment with alkaline phosphatase (AP, black arrow in C). In **B** and **C**, the position of CotB-66 and CotG-36 (blue arrows), CotB-34 (grey arrow), CotB-46 (red arrow), CotH or CotH^D228Q^ (green arrows), high molecular weight forms of CotG (black arrows in C, bottom panel), is indicated. The position of molecular weight markers (MW, in kDa) is shown on the left side of the two panels.

The coat protein profile of a *cotH* insertional mutant was originally established after SDS/DTT extraction of the spore coat proteins (Naclerio *et al.*, 1996). At least 9 proteins are absent from spores of a *cotH* insertional mutant, including CotG-36 and CotB-66, CotS (41 kDa), CotSA (42.9 kDa), CotQ (50 kDa), CotU (11.6 kDa) and CotC (14.8 kDa), YusA (30.4 kDa) and CotZ (16.5 kDa; but see below) (Baccigalupi *et al.*, 2004, Giorno *et al.*, 2007, Kim *et al.*, 2006, Naclerio *et al.*, 1996, Zilhao *et al.*, 1999, Zilhao *et al.*, 2004) (NB: hereafter, the number following the designation of a particular coat protein will refer to its apparent molecular weight under SDS-PAGE conditions). Here, spores were density gradient purified and the coat proteins were extracted with NaOH and analyzed by SDS-PAGE and immunoblotting (Henriques & Moran, 2007). NaOH extraction was used because in preliminary work, we failed to detect CotH by immunoblotting in SDS/DTT coat extracts; presumably not sufficient protein is extracted by this method to allow detection by our antibody. For reference, however, an SDS-PAGE analysis of coat extracts obtained from *cotH* and *cotH*^*D228Q*^ spores by SDS/DTT extraction is shown in Figure S2A (see also the Supplemental text). The collection of coat proteins obtained from *cotH* spores following NaOH extraction revealed the absence of CotB-66, CotG-36, and CotC and at least four other proteins in the 30-50 kDa region of the gel (Fig. 2B, red and blue arrows in the top panel). In addition, at least three proteins were more extractable from *cotH* or *cotH*^*D228Q*^ spores (Fig. 2B, brown arrows). Importantly, the pattern of extractable coat proteins from *cotH* or *cotH*^*D228Q*^ spores was nearly identical, while insertion of the WT *cotH* allele at *amyE* restored the WT pattern to either mutant, as assessed by Coomassie staining (Fig. 2B, top). Spores of a *cotH* insertional mutant as well as those of the catalytically inactive *cotHD251A* are impaired in L-alanine-triggered spore germination (Naclerio *et al.*, 1996, Nguyen *et al.*, 2016). A mixture of L-Asp, glucose, fructose and KCl (AGFK) activates a second pathway of nutrient-induced spore germination (Setlow, 2014). We show that *cotH* or *cotHD228Q* spores are impaired in AGFK-triggered spore germination and that complementation with the WT allele restores normal germination (Fig. S2B). Thus, the kinase activity of CotH influences the two known pathways of nutrient-induced spore germination (Setlow, 2014, Setlow *et al.*, 2017).

Consistent with previous results (Giglio *et al.*, 2011), immunoblot analysis of coat extracts prepared from purified spores shows that the level of coat-associated CotH^WT^ produced from the *amyE* locus under the control of the native *cotH* promoter was similar to that found for a WT strain (Fig. 2B, bottom panel). Importantly, the level of coat-associated CotH^D228Q^ was only slightly lower that of the WT protein, indicating that the activity of CotH is not required for its own assembly (Fig. 2B). In contrast, the catalytic inactive CotH^D251^ was not incorporated into the coat, unless overproduced from stronger promoters (Nguyen *et al.*, 2016). Therefore, the *cotHD228Q* mutant may more accurately reflect the impact of the absence of active CotH on the overall assembly and properties of the spore coat. While it seems plausible that the assembly of the CotH-controlled proteins requires their phosphorylation by CotH, the possibility that CotH acts as a priming kinase for another, as yet unknown kinase has been raised and cannot presently be excluded (Nguyen *et al.*, 2016).

### CotG is required for the phosphorylation of CotB-46 in spores

CotH catalyzed incorporation of ^32^P from [γ-^32^P]-ATP into purified CotB-46, shifting its SDS-PAGE mobility to the 66 kDa region of the gel (Nguyen *et al.*, 2016). Thus, in the spore coat, CotB-66 is also likely to arise through extensive phosphorylation of CotB-46, an inference in line with the detection of a phosphorylated species of about 66 kDa in spores of strains producing CotH^WT^ but not CotH^D228Q^ (Nguyen *et al.*, 2016). A phosphorylated species inferred to be CotG was also detected (Nguyen *et al.*, 2016). In this study, we wanted to examine extracts of a *cotG* mutant, since CotG is known to be required for formation of CotB-66 in spores (Naclerio *et al.*, 1996, Zilhao *et al.*, 2004), whereas in the study of Nguyen and co-authors CotB-66 could be formed through direct phosphorylation of purified CotB-46 by CotH, that is, in the absence of CotG (Nguyen *et al.*, 2016). Coat protein extracts were prepared from purified spores of several strains by NaOH treatment, analyzed by SDS-PAGE and Coomassie staining and additionally by immunobloting with anti-phosphoserine (pSer), anti-CotB, and anti-CotG antibodies before and after incubation with alkaline phosphatase. CotB-66 was detected in Coomassie-stained gels of WT extracts but was absent from *cotG, cotH* (in these two mutants only CotB-46 is detected) and *cotB* spore coat extracts (Naclerio *et al.*, 1996, Zilhao *et al.*, 2004) (Fig. 2C, top panel). CotB-66 reacted with the anti-pSer and anti-CotB antibodies (Fig. 2C, two middle panels). A form of CotB, CotB-34, detected by immunoblotting in all the extracts except in those from *cotB* and *cotH* mutants, did not react with anti-pSer (Fig. 2C, black arrow in the second panel from the top); it may correspond to a proteolytic fragment encompassing the N-terminal moiety of the protein (expected size of about 28.7 kDa; see also below). Treatment of the WT spore coat extracts with calf intestinal phosphatase (AP) caused disappearance of CotB-66 from the Coomassie-stained gels and from the immunoblots in which anti-CotB or anti-pSer antibodies were used (Fig. 2C). Concurrently, AP treatment resulted in the appearance of CotB-46, which reacted with the anti-CotB antibody but not with anti-pSer (Fig. 2C). The presence of CotB-66 was restored in the *cotH* complementation strain and treatment with AP strongly reduced its level; because the level of CotB-66 is higher in this strain than in the WT, however, the AP treatment did not completely convert CotB-66 into CotB-46. Rather, it resulted in a smear corresponding to several bands below CotB-66, which reacted both with anti-CotB and anti-pSer antibodies (Fig. 2C). No form of CotB reacted with the anti-pSer or the anti-CotB antibodies in the coat extract prepared from *cotH* or *cotHD228Q* spores (Fig. 2C). Furthermore, in the *cotG* mutant only CotB-46 form is detected with the anti-CotB antibody and no form of CotB reacted with the anti-pSer antibody (Fig. 2C). Together, these results indicate that in the coat, CotB-46 is not phosphorylated (or its phosphorylation is below our detection level), and that CotB-66 is formed from CotB-46 through CotH-mediated phosphorylation.

In the WT, CotG is detected in coat extracts as a diffuse band around 36 kDa (CotG-36) when extracts are prepared by SDS/DTT treatment ((Naclerio *et al.*, 1996, Zilhao *et al.*, 2004); see also Fig. S2A). Following NaOH treatment CotG-36 is only weakly detected by Coomassie staining (Fig. 2B and C, top) but immunobloting with an anti-CotG antibody reveals its presence in the extracts (Fig. 2C, bottom panel). CotG-36 was also not detected by the anti-pSer antibody (Fig. 2C). Nevertheless, CotG-36 may correspond to a phosphorylated form of CotG because it is no longer detected by immunoblotting following AP treatment of the extracts; instead, several bands are detected below CotG-36 (Fig. 2C; see also below). Also, in *cotB* spores and also in the *cotH* insertional mutant complemented with WT *cotH* at *amyE*, at least one additional anti-CotG-reactive band is detected above CotG-36 (Fig. 2C, black arrows in the bottom panel). Detection of an additional CotG form above CotG-36 in spores of the *cotB* mutant raises the possibility that phosphorylation of CotG is enhanced in the absence of CotB. In line with earlier work (Zilhao *et al.*, 2004), no CotG is detected by immunoblotting in *cotH* or *cotHD228Q* spores (Fig. 2C).

In all, the results indicate that CotG-36 and CotB-66, the two main forms of the proteins detected in spores, are phosphorylated in a CotH-dependent manner, in agreement with the identification of phosphosites in both CotG and CotB and the identification of CotB-46 and a CotG-derived peptide as direct substrates of CotH ((Nguyen *et al.*, 2016, Rosenberg *et al.*, 2015, Saggese *et al.*, 2014); this work). While un-phosphorylated CotB-46 associates with spores, un-phosphorylated CotG is unstable, consistent with the difficulty in overproducing the protein in *E. coli* ((Nguyen *et al.*, 2016); see also below).

### CotG is necessary and sufficient for the efficient phosphorylation of CotB-46 by CotH in *E. coli*

Phosphorylation of purified CotB-46 by CotH *in vitro* in the presence of [γ-32P]ATP appeared inefficient, as a clear autoradiography signal was only detected after an incubation of 2 h (Nguyen *et al.*, 2016). Since CotG is required for phosphorylation of CotB-46 in spores, we considered the possibility that CotG could somehow promote the phosphorylation of CotB-46. We overproduced CotB-46 in *E. coli* cells through an auto-induction regime, alone or together with CotG and/or CotH^WT^ or CotH^D228Q^ (Fig. 3). Proteins in lysates prepared from the various strains were resolved by SDS/PAGE and the phosphorylation status of CotB and CotG assessed by Coomassie staining and immunoblotting, using an anti-pSer antibody. CotH^WT^ and CotH^D228Q^ both accumulated as species of about 40 kDa, consistent with their calculated mass (about 42.8 kDa; Fig. 3). In agreement with previous results (Nguyen *et al.*, 2016, Zilhao *et al.*, 2004), expression of *cotB* alone resulted in the accumulation of CotB-46, which did not react with anti-pSer (Fig. 3B) and did not stain with the Pro-Q diamond dye (Fig. S3; see also the supplemental text). CotB-46 also accumulated when *cotB* and *cotG* were co-induced together with *cotHD288D* (Fig. 3B). Strikingly, however, the co-induction of *cotB* with *cotH* and *cotG* resulted in the complete conversion of CotB-46 into CotB-66 (Fig. 3A). CotB-66 reacted with both the anti-CotB and the anti-pSer antibodies (Fig. 3A and B, bottom) and was stained by the Pro-Q diamond dye (Fig. S3).

**Figure 3.**
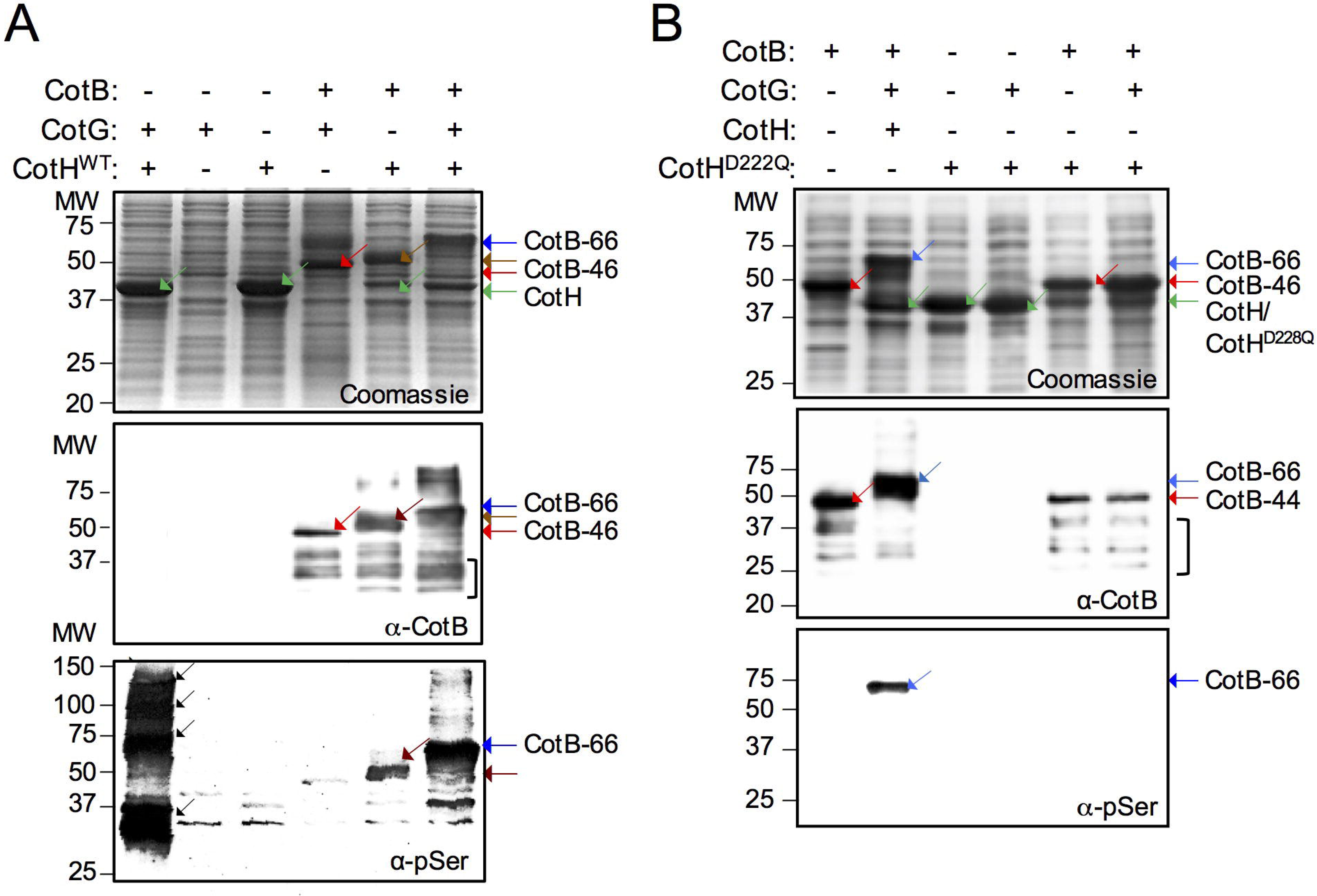
CotG is necessary and sufficient for the efficient phosphorylation of CotB-46 by CotH. CotH^WT^ (**A**) or CotH^D228Q^ (**B**) were overproduced from a T7*lac* promoter in *E. coli* strains alone or together with the indicated proteins (“+” signs). The strains were grown in autoinduction medium, whole cell extracts prepared and the proteins resolved by SDS-PAGE. Coomassie stained gels (top) and the immunoblot analysis with anti-CotB (middle panels) or anti-pSer antibodies (bottom) are show for both **A** and **B**. The position of relevant species is indicated on the right side of the panel. The parentheses in **A** and **B** (middle panels) indicates possible degradation forms of CotB. See also Fig. S3, which shows the same gels stained with Sypro Ruby or Pro-Q Diamond. In panels **A** and **B**: the blue arrow shows the position of CotB-66, and the red arrow the position of CotB-46; a form of CotB with a mobility between that of CotB-46 and CotB-66 is indicated by a brown arrow; CotH^WT^ (or CotH^D228Q^) is indicated by a green arrow. The position of molecular weight markers is shown on the left side of all panels.

A form of CotB above CotB-46 but below CotB-66 accumulates when only *cotB* and *cotH* are co-induced (Fig. 3A and Fig. S3A); this species reacts with the anti-pSer antibody but presumably it is not as extensively phosphorylated as CotB-66, explaining its intermediate mobility. CotG did not accumulate, as assessed by Coomassie staining, when produced alone or in combination with CotB and/or CotH (Fig. 3). The co-induction of *cotG* and *cotH*, however, resulted in the formation of species with apparent mobilities around 50, 75, 100 and 150 kDa, which were detected by the anti-pSer antibody (Fig. 3A). This pattern of phosphorylated species is different from the one obtained upon co-induction of *cotB, cotG* and *cotH* (Fig. 3A), suggesting that CotG may only be efficiently phosphorylated in the absence of CotB. Conversely, although CotH can phosphorylate CotB-46 as also shown previously (Nguyen *et al.*, 2016), efficient phosphorylation of the protein requires CotG and no additional factor.

### Direct phosphorylation of CotG by CotH

In an attempt to determine whether full-length CotG could be phosphorylated by CotH, at least in the absence of CotB, we first overproduced CotH^WT^ or CotH^D228Q^ with a C-terminal *Strep*-tag in *E. coli*, and the two proteins were affinity purified (see Material and Methods). The presence of the *Strep-*tag did not impair kinase activity, as shown by the formation of CotB-66 when *cotH* was induced together with *cotB* and *cotG* (Fig. S4). Purified CotH^WT^ but not CotH^D228Q^ showed auto-phosphorylation activity in the presence of [γ-^32^P]ATP (Nguyen *et al.*, 2016) (Fig. S5A). Moreover, CotH^WT^ but not CotH^D228Q^ showed trans-phophorylation activity when incubated with purified CotB-46 in the presence of [γ-^32^P]ATP (Fig. S5B). Although labeling of CotB-46 was detected, formation of CotB-66 was not presumably because under our experimental conditions the degree of phosphorylation of CotB-46 in the absence of CotG is insufficient to alter its electrophoretic mobility ((Nguyen *et al.*, 2016); Fig. S5B). Labeled CotH^WT^ was only detected in the absence of CotB-46, suggesting rapid transfer of phosphoryl groups to CotB-46. Staurosporine (STP), a small-molecule ATP mimic, is a powerful inhibitor of some intracellular eukaryotic Ser/Thr kinases while others, atypical, are insensitive to it (Ruegg & Burgess, 1989, Xiao *et al.*, 2013). We found that both the auto- and trans-phosphorylation activity of CotH were insensitive to STP at concentrations up to 70 μM (Fig. S5). In contrast, at a concentration as low as 10 µM, STP inhibited the PrkC-dependent germination of spores in response to peptidoglycan fragments (Shah *et al.*, 2008). While in most kinases the adenine part of ATP is located within a pocket of hydrophobic residues, the structure of CotH in a complex with Mg^2+^/AMP shows the adenine moiety of AMP sandwiched between two aromatic residues Tyr142 and Trp245 (Nguyen *et al.*, 2016). Presumably, the bulky side chains of these residues blocks binding of the inhibitor to the active site (Xiao *et al.*, 2013).

To test whether CotH could directly phosphorylate CotG, we used immunoblotting to monitor the accumulation of CotG over time following IPTG induction of *cotG* alone or upon co-induction of *cotH* (Fig. 4). Under these conditions, CotG was initially detected as a species of about 34 kDa (Fig. 4, red arrow in the two panels) that may correspond to the un-phosphorylated form of the protein. Over time, in the presence of CotH^WT^ (but not in the presence of CotH^D228Q^, CotG-36 accumulated (Fig. 4, blue arrow). Additional species, with an apparent molecular weight greater than 36 kDa, were also detected upon prolonged incubation with CotH^WT^ (Fig. 4B, black arrows). Importantly, formation of all the forms of CotG above CotG-34 decreased when *cotB* was co-induced (Fig. 4B). We conclude that full-length CotG is directly phosphorylated by CotH. Moreover, our results suggest that phosphorylation of CotG is less efficient in the presence of CotB.

**Figure 4.**
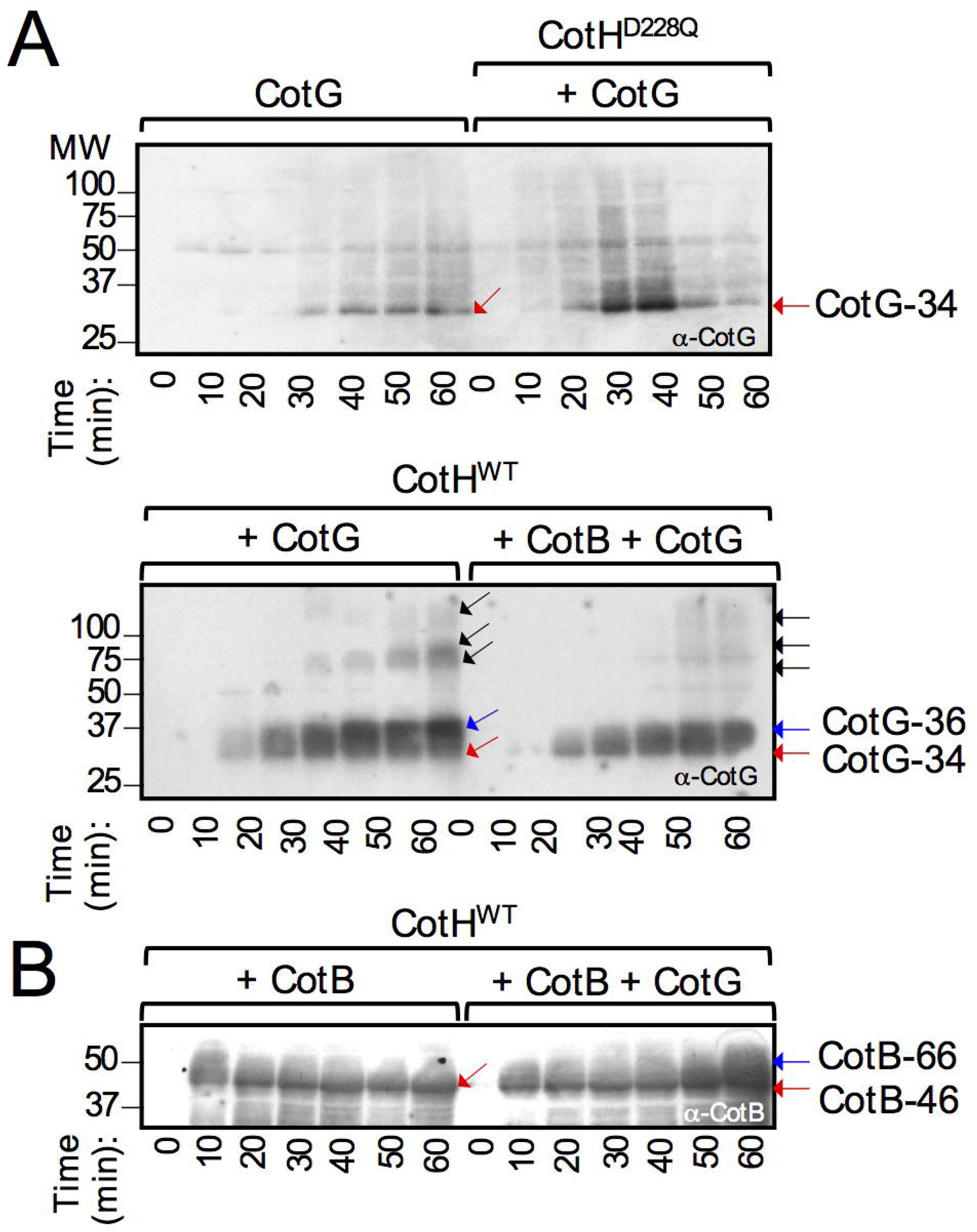
Direct phosphorylation of CotG by CotH. IPTG was added to *E. coli* strains producing either CotG alone or in the presence of CotH^D228Q^ (A), or CotG with CotH^WT^ in the absence (B, left) or in the presence (B, right) of CotB. The proteins were produced from a T7*lac* promoter and samples were taken from the cultures at the indicated times (in min) after IPTG addition. The samples were subject to immmunoblot analysis with an anti-CotG antibody. The position of CotG-34 is shown by a red arrow and CotG-36 is indicated by a blue arrow. The position of forms of CotG-34 with apparent masses higher than that of CotG-36 is shown by black arrows. Molecular weight markers (in kDa) are shown on the left side of the panels.

### CotG interacts with the C-terminal region of CotB

We have shown before that CotB and CotG self-interact and that CotB interacts with CotG (Zilhao *et al.*, 2004). That the SKR^B^ region is the site of CotB-46 phosphorylation, together with the requirement for CotG for the formation of CotB-66, suggested to us that CotG could specifically interact with the C-terminal moiety of CotB. To test this possibility, we used a GAL4-based yeast two-hybrid interaction assay (Zilhao *et al.*, 2004). We fused the entire coding sequence of CotB (CotB^FL^) or CotG to either the activation (AD) or the DNA-binding domain (BD) of GAL4 yeast transcriptional activator GAL4 (Table 1). In addition, the sequences coding for the N-(CotB^N^, residues 1-213) or C-terminal (CotB^C^,residues 195-380) regions of CotB^FL^ were also fused to both the AD and BD of GAL4. Interactions between the various fusion proteins were assessed in a yeast reporter strain using a colony lift assay that monitors expression of the *lacZ* gene (Zilhao *et al.*, 2004). Under our experimental conditions, background levels of β-galactosidase activity, determined by co-transforming the same cells with the two empty vectors, were negligible (Table 1). Also, no β-galactosidase activity was detected when individual fusion proteins were expressed with the corresponding empty vector control. We found an interaction between CotB^FL^ and itself and a stronger interaction of CotB^N^ with itself (Table 1). This suggests that the SKR^B^ region of CotB somehow reduces the ability of the N-terminal moiety to self-interact. In contrast, we found no evidence for an interaction of CotB^C^ with itself. Both CotB^FL^ and CotB^C^ interacted with CotG, but CotB^N^ did not, nor did a form of CotB deleted for the SKR^B^ region (Table 1). Thus, it is the C-terminal moiety of CotB that interacts with CotG, but only if the SKR^B^ region is present. The results suggest that CotB interacts with CotG via the SKR^B^ region, and it seems possible that this interaction is part of the mechanism by which CotG promotes phosphorylation of CotB-46. It also seems possible that this interaction somehow reduces the ability of CotH to phosphorylate CotG.

**Table 1.** Detection of *lac*Z transcription by colony lift assays in yeast diploid strains Y187/Y190 containing fusions of products encoded by *cot*G, *cotH*, and full length, N-or C-terminal regions of *cotB* to GAL4 activation and binding domains.

| 1000 | DNA-binding domain fusion<sup>b</sup> |  |  |  |  |  |  |
| --- | --- | --- | --- | --- | --- | --- | --- |
| 1001 |  |  |  |  |  |  |  |
| 1002 | Activation domain | pAS2-1 | pAS-B | pAS-BN | pAS-BC | pAS-G | pAS-H |
| 1003 | fusion<sup>a</sup> | (BD) | (CotB-FL-BD) | (CotB-N-BD) | (CotB-C-BD) | (CotG-BD) | (CotH-BD) |
| 1004 |  |  |  |  |  |  |  |
| 1005 | pACT-2 (AD) | -<sup>c</sup> | - | - | - | - | - |
| 1006 | pAC-B/(CotB-FL-AD) | - | + | + | - | + | - |
| 1007 | pAC-BN/(CotB-N-AD) | - | + | +++ | - | - | - |
| 1008 | pAC-BC /(CotB-C-AD) | - |  | - | - | + | - |
| 1009 | pAC-G/ (CotG-AD) | - | + | - | + | + | - |
| 1010 | pAC-H/(CotH-AD) | - | - | - | - | - | - |
| 1011 |  |  |  |  |  |  |  |

<sup>a</sup> Description of plasmid constructs in pACT-2 (contains GAL4 activation domain (AD)) that were transformed into yeast strain Y190 (Clontech).
<sup>b</sup> Description of plasmid constructs in pAS2-1 (binding domain (BD)) that were transformed into yeast strain Y187 (Clontech).
<sup>c</sup> (-/+) represent the time for detection of blue colonies on a colony lift assay for *lacZ* expression; +++, +, and - indicates development of color in 30 minutes, 1hour, and greater than one hour, respectively, as determined by testing 3 independent colonies for each pairwise combination.

### The kinase activity of CotH patterns the spore outer coat

Deletion of *cotH* brings about drastic alterations on the ultrastructure of the spore outer coat as viewed by transmission electron microscopy (TEM) (Zilhao *et al.*, 1999). We examined spores of the *cotH* and *cotHD228Q* mutants along with spores of the congenic WT strains, by TEM. In both *cotH* and *cotHD228Q* mutants, and although the region normally occupied by the outer coat had approximately the same width as in WT spores, the electrondense outer coat striations were absent and replaced by partially structured material (Fig. 5). The outer edge of the coat, however, showed a defined contour. In addition, the inner coat lamellae appeared reduced in both mutants (Fig. 5).

**Figure 5.**
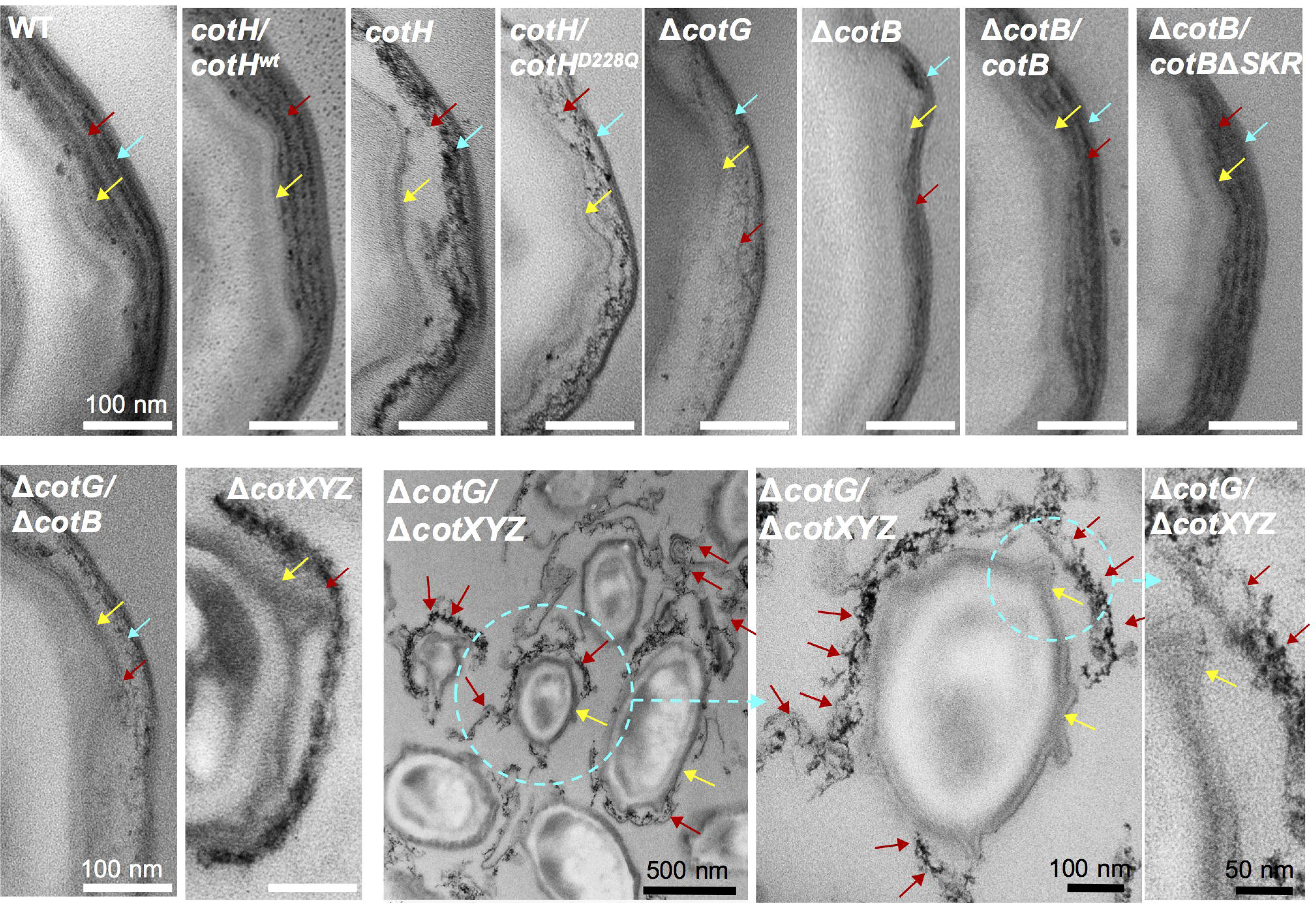
CotH patterns the spore outer coat. **A:** Transmission electron microscopy of spores produced by the following strains, as indicated: wild type (WT), *cotH*/*cotH*^*wt*^, *cotH, cotH*/*cotH*^*D228Q*^, δ*cotG,* δ*cotB*, and δ*cotB* complemented *in trans* with either WT *cotB* or *cotB*δSKR, δ*cotXYZ*, and δ*cotG/* δ*cotXYZ.* Spores were collected from DSM cultures 24 h after the initiation of sporulation. The two bottom panels show a field of δ*cotG/* δ*cotXYZ* spores (left) and higher magnification images (two last panels on the right) of the region encircled. The brown arrow points to the outer coat or electrondense outer coat material, the yellow arrow to the inner coat, and the blue arrow to the crust region. The green arrows in the δ*cotXYZ* panel point to discontinuities in the outer coat layer.

The structure of the coat in the *cotH* or *cotHD228Q* mutants is reminiscent of that reported for spores of a *cotG* insertional mutant, in which the electrondense outer coat striations are replaced by partially structured material, but the outer edge of the coat remains delimited by a thin well-defined layer (Henriques *et al.*, 1998). Thus, *cotG* was proposed to be an important structural organizer of the spore outer coat. As already mentioned, however, the *cotH* promoter is now known to be located downstream of the 3′-end of *cotG* and *cotG* insertional mutations predictively exert a polar effect on *cotH* expression (Giglio *et al.*, 2011) (Fig. 1A). We therefore constructed a strain bearing an in-frame *cotG* deletion and characterized the spores formed by this new mutant by TEM. We found that spores of the new *cotG* deletion mutant to be essentially indistinguishable from those of the insertional mutant used before (Henriques *et al.*, 1998) (Fig. 5). Thus, this result not only supports the original conclusion that CotG is a key organizer of the spore outer coat (Henriques *et al.*, 1998) but further suggests that the kinase activity of CotH is required for proper formation of the spore outer coat, via *cotG*. Possibly, the phosphorylation of CotG stabilizes the protein while allowing it to promote formation of the organizational pattern normally seen for the outer coat. Both the phosphorylation of CotG and CotB-46 are likely to occur mostly at the spore surface (Fig. S6; see also the supplemental text).

### The SKR^B^ region is dispensable for proper coat morphogenesis

Because CotB-66 is not formed in *cotG* mutants, we reasoned that the CotH-dependent phosphorylation of CotB-46, promoted by CotG, could also be a key determinant for the structural organization of the outer coat. If so, then spores of a *cotB* mutant could share some of the structural features observed for *cotH* or *cotG* spores. To test this, we first constructed a *cotB* in-frame deletion mutant and examined the spores produced by this strain by TEM. We found that spores of a Δ*cotB* mutant had a thinner outer coat but that the pattern of electrondense striations was retained (Fig. 5, blue arrow). This suggests that phosphorylation of CotB-46 is not a main determinant of the structural organization of the outer coat.

To further test this inference, we constructed strains bearing an in-frame deletion of the *cotB* gene and either a WT *cotB* allele or an allele with an in-frame deletion of the SKR^B^ region at *amyE* (Fig. S7A and B). Analysis of the proteins extractable from the coat of Δ*cotB/cotBWT* and Δ*cotB*/*cotBΔSKR* spores shows a similar collection of proteins except for the absence of CotB-66 from spores of the latter strain, an observation confirmed by Immunoblot analysis (Fig. S7C). Thus, as previously reported for a *cotB* insertional allele (Naclerio *et al.*, 1996, Zilhao *et al.*, 2004), neither the Δ*cotB* nor the *cotBΔSKR* alleles cause gross alterations in the composition of the coat.

Spores of the Δ*cotB* mutant with either WT *cotB* or *cotBΔSKR* in trans were then analyzed by TEM (Fig. 5). Both showed an electrondense striated outer coat and appeared indistinguishable from WT spores (Fig. 5). Thus, the SKR^B^ region and hence phosphorylation of CotB-46, is not a prerequisite for the structural organization of the outer coat. By comparison with the thinner outer coat of Δ*cotB* spores, it follows that the CotB NTD contributes to the normal thickness of the spore outer coat.

### CotH activity is not essential for crust assembly

A thin, well-defined layer forms the outer edge of the coat in *cotH*, Δ*cotB* or Δ*cotG* mutants ((Henriques *et al.*, 1998, Zilhao *et al.*, 1999); Fig. 5). This structure could correspond to the crust, or a crust basal-layer. Assembly of the crust is dependent on the outer coat morphogenetic protein CotE (McKenney *et al.*, 2010) which controls CotH assembly (Isticato *et al.*, 2015, Naclerio *et al.*, 1996, Zilhao *et al.*, 1999). Yet, at least the crust protein CotW is independent on CotH for assembly (Kim *et al.*, 2006). To test whether CotH was required for assembly of the crust, and since CotZ is required for crust assembly (McKenney *et al.*, 2010), we examined the localization of a CotZ-GFP fusion in spores of the WT and in spores of the *cotH*, Δ*cotG* and Δ*cotB* mutants. We found that CotZ-GFP formed a complete ring of fluorescence around 62% of the WT spores scored, and a polar cap in 38% of the spores (Fig. 6). These numbers did not differ much for the *cotHD228Q*, Δ*cotB*, Δ*cotG* or *cotH* spores, suggesting that in these mutants the crust is still assembled. Nevertheless, in spores of the strain bearing the WT *cotH* allele at *amyE*, the cap pattern of CotZ-GFP fluorescence was reduced to 13%, while the complete circle pattern increased to 87% (Fig. 6). Thus, while largely dispensable for crust assembly, CotH may somehow influence formation of this structure.

**Figure 6.**
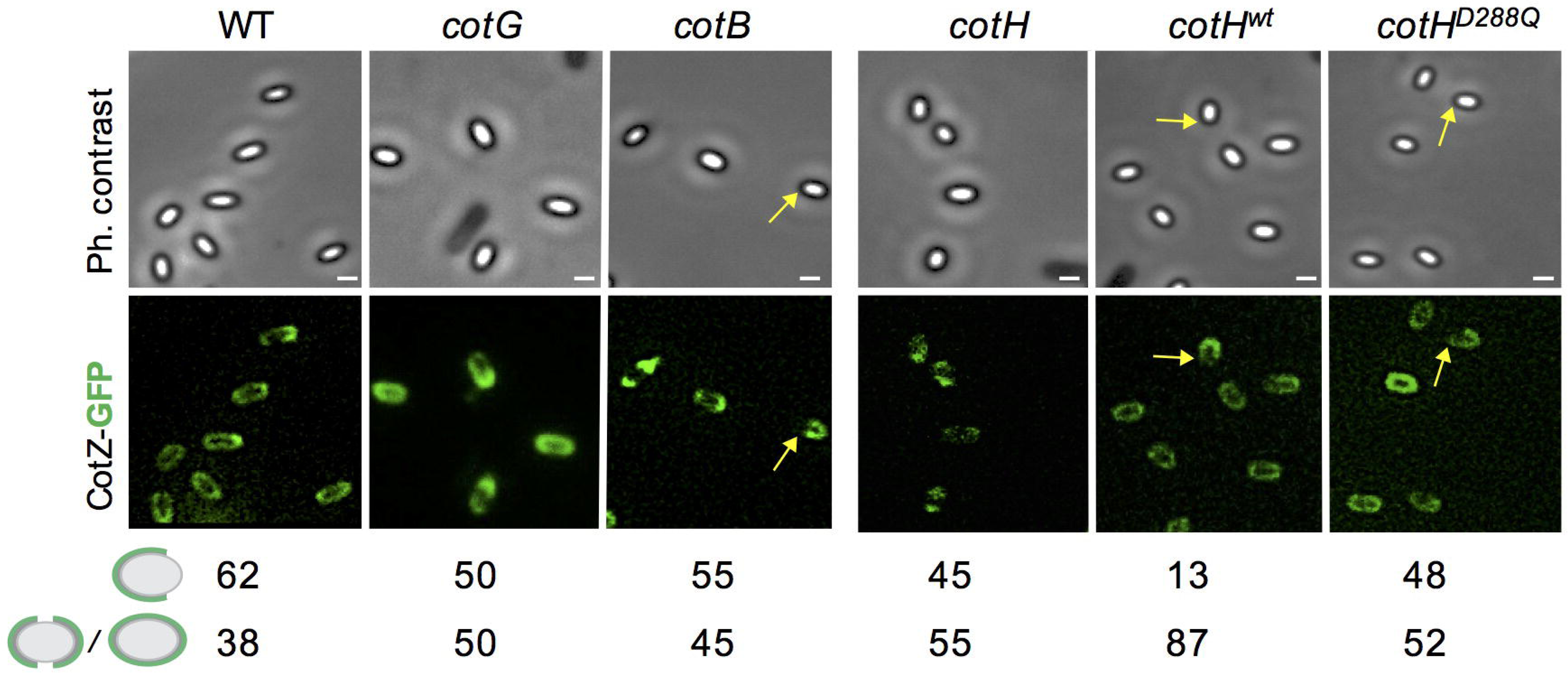
Localization of CotZ-GFP in mature spores. Spores of the indicated mutants expressing a *cotZ-GFP* translational fusion were purified and analysed for the localization of the fusion protein. The percentage of the two main patterns of fluorescence observed for CotZ-GFP, full circle and one or two polar caps (a single cap is represented for simplicity), relative to the total number of spores scored (*n*) is indicated for each strain. Scale bar, 0.2 µm.

We introduced a deletion of the *cotX, cotY* and *cotZ* genes (Δ*cotXYZ*) (Zhang *et al.*, 1993) into the Δ*cotG* mutant, as an independent test of the idea that the thin structure seen at the edge of Δ*cotG* spores corresponds to the crust. As a control, we also examined spores of the Δ*cotXYZ* mutant. As previously reported (Zhang *et al.*, 1993), the outer coat of Δ*cotXYZ* spores is less organized and shows discontinuities around the periphery of the spores (Fig. 5). Consistent with the view that the crust forms the edge of the outer coat region in Δ*cotG* spores, this layer is absent from spores of the Δ*cotG*/Δ*cotXYZ* mutant. Strikingly, the spores are often surrounded by detached electrondense material that often projects into the surrounding medium forming long twirls (Fig. 5). We posit that this material likely to correspond to the patches of electrondense amorphous material seen in the outer coat region of Δ*cotG* spores.

The phenotype of Δ*cotXYZ* and Δ*cotG*/Δ*cotXYZ* spores suggests that the crust has a role in maintaining the integrity and localization of the outer coat or outer coat material. The data are also consistent with a model in which *cotG* and *cotH* are essential determinants of the normal patterning of the outer coat, that the kinase activity of CotH is required mainly for outer coat assembly, and that the normal structural organization of the outer coat is not an essential pre-requisite for crust assembly.

## Discussion

CotH is a eukaryotic-type Ser/Thr kinase responsible for the phosphorylation of at least 28 serine residues within the SKR^B^ region of CotB-46 and at least 15-16 residues in the SKR^G^ region of CotG. Moreover, we show for the first time that CotB-46 is phosphorylated multiple times per molecule. It is the extensive phosphorylation of CotB-46 that slows its mobility under SDS-PAGE conditions causing the protein to migrate with an apparent mass of 66 kDa. Evidence also suggests that the phosphorylation of CotG results in the formation of CotG-36, the main form of the protein found in the coat by Coomassie staining, as well as other forms of CotG, of higher apparent mass, that are also detected in the coat by immunoblotting ((Zilhao *et al.*, 2005); this work). We have used the catalytically inactive CotH^D228Q^ protein, to examine the overall role of CotH in coat assembly and structure. Another catalytically inactive form of CotH, CotH^D251A^, has been analysed for the presence of phosphorylated proteins in the coat by immunoblotting, but gels documenting the collection of proteins extracted from spores of the mutants have not been reported (Nguyen *et al.*, 2016). CotH^D228Q^ results in spores which lack the same proteins absent from a *cotH* insertional mutant. Importantly, while CotH^D228Q^ is assembled into the spore coat under the control of its normal promoter, CotH^D251A^, was only assembled when overproduced from the P_*cotA*_ or P_*gerE*_ promoters (both σ^K^ dependent and GerE-repressed) or the P_*cotE* *P1*_ promoter (σ^K^-controlled) (Nguyen *et al.*, 2016). Altering the time and level of *cot* gene expression, however, may impact drastically on assembly of the coat (see for example, (Costa *et al.*, 2007)). Moreover, overexpression of *cotH* bypasses the need for CotE for the assembly of several CotE-dependent proteins (Isticato *et al.*, 2013). Thus, CotH^D228Q^ reveals that the absence of kinase activity and most likely no other perturbation in the assembly pathway independently of kinase activity, has a global impact on coat assembly. The proteins missing in *cotH228* spores include CotB-66, CotG and CotC, as well as other proteins presumed to be CotH-controlled and that may also be phosphorylated by CotH. Several coat proteins phosphorylated at Ser residues, including the CotH-controlled proteins CotO, and YhcQ, YqfQ, GerW, and YtxO, have been detected during spore germination (Rosenberg *et al.*, 2015). Spores of a *cotHD251A* mutant are also impaired in L-Ala-triggered germination (Nguyen *et al.*, 2016), even if part of the effect could result from secondary effects of the overexpressed allele on coat assembly. In any event, and since *cotHD228Q* spores are impaired in AGFK-triggered germination, the kinase activity of CotH directly affects the two pathways known to control the germination of spores in response to nutrients (Setlow, 2014).

Co-production of CotB and CotH in *E. coli* is not sufficient to shift migration of the protein to the 66 kDa region of the gel; in the presence of CotG, however, CotB-66 is promptly and efficiently formed (Fig. 3). Conversely, the CotH-dependent phosphorylation of CotG is enhanced in *E. coli*, in the absence of CotB-46. These reactions mimic the situation during coat assembly (Fig. 7A). In the absence of CotG, CotB-46 may be phosphorylated, but not sufficiently to alter its SDS-PAGE migration noticeably (Fig. 7A). In contrast, in the absence of CotB, phosphorylation of CotG seems more extensive (Fig. 7A). How CotG stimulates the phosphorylation of CotB-46 is unknown. Phosphorylation of CotG, however, is not a pre-requisite for the interaction with and the phosphorylation of CotB-46 by CotH, because CotG^SKR^ still promotes formation of CotB-66 *in vivo*, in a CotH-dependent manner (Saggese *et al.*, 2016) and because the interaction between CotG and CotB is detected in yeast cells in the absence of CotH (Table 1). CotG binds to the C-terminal moiety of CotB (Table 1) and it seems plausible that this interaction involves the N- and C-terminal regions of CotG flanking the SKR^G^ region (Saggese *et al.*, 2016) (Fig. 7B). Possibly, binding of CotG to CotB alters the conformation of and/or exposes the SKR^B^ region, predicted to be disordered, facilitating phosphorylation by CotH (Fig. 7B). In contrast, the interaction of CotG with CotB may impair phosphorylation of CotG by CotH.

**Figure 7.**
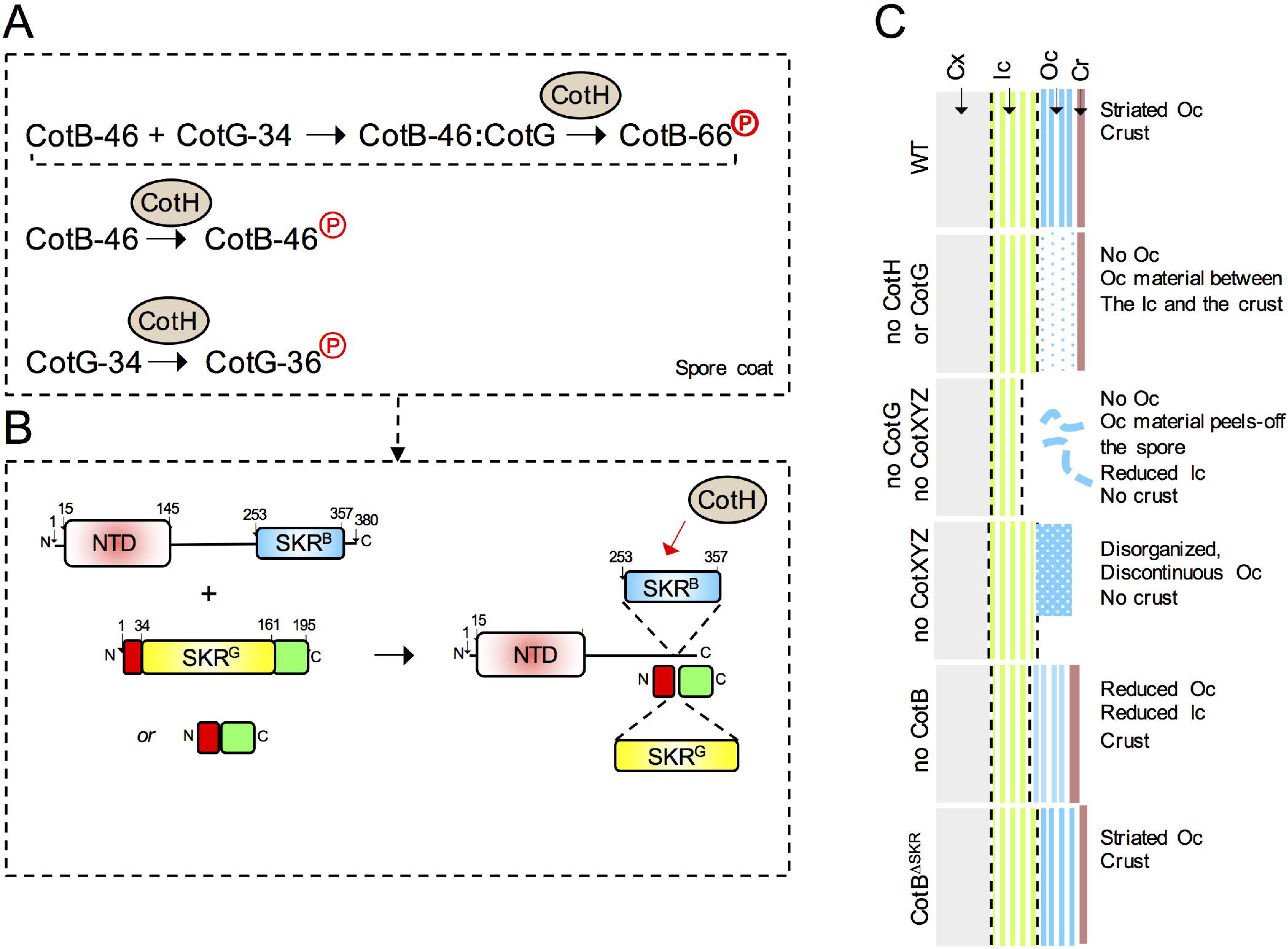
**A**: CotH-dependent phosphorylation reactions. CotG-34 interacts with CotB-46 to form a complex that allows the phosphorylation of the latter to form CotB-66 (phosphoryl group, P, in bold). CotH can also direct the low level phosphorylation (phosphoryl group, P) of CotB-46 in the absence of CotG, and the phosphorylation of CotG-34. All three proteins are assembled onto the spore outer coat, where the represented reactions occur. CotH is likely to phosphorylate other, as yet unidentified, spore coat proteins. **B**: CotG interacts with the C-terminal moiety of CotB-46. The complex is though to expose the SKR^G^ region, facilitating phosphorylation by CotH (red arrow). It is possible that the SKR^G^ region of CotG is also phosphorylated at this stage. The interaction with CotB is likely to involve regions outside of SKR^G^, because a form of CotG with a deletion of the SKR^G^ region still promotes formation of CotB-66 *in vivo*. **C**: Schematic representation of the spore surface layers in the WT and in various mutants. In the WT, the outer coat (Oc, blue) has an electrondense striated pattern and the crust is closely apposed (Cr, brown). In *cotH* and *cotG* mutants, the electrondense striated OC is absent, replaced by a space where amorphous material is seen, delimited by the crust. δ*cotB* and *cotB*δ*SKR* spores show an adherent crust, but δ*cotB* spores may have a thinner, less electrondense outer coat. The reduced inner coat (Ic, green) in the mutants may be an indirect effect of improper formation of the outer coat. Cx, spore cortex.

The SKR^G^ region is also predicted to be disordered and may be susceptible to proteolysis (Giglio *et al.*, 2011, Saggese *et al.*, 2016). The repeat region of *B. anthracis* ExsB, a CotG homologue, was also found to be extremely sensitive to proteolysis (McPherson *et al.*, 2010). Phosphorylation may thus stabilize the protein, explaining why CotG does not accumulate in spores unable to produce active CotH or in *E. coli* ((Isticato *et al.*, 2015, Zilhao *et al.*, 2004); this work). Previous work has shown that CotH stabilizes two other outer coat proteins, CotC and CotU, in the mother cell when their assembly is prevented by deletion of *cotE* (Isticato *et al.*, 2004, Isticato *et al.*, 2008, Isticato *et al.*, 2013). Thus, CotH-mediated phosphorylation may serve a general role in the stabilization of coat proteins following their synthesis in the mother cell, and in promoting the formation of protein complexes competent for assembly (Isticato *et al.*, 2013, Isticato *et al.*, 2015). CotH, however, is also active at the spore surface where, over time, CotB-46 is converted into CotB-66 and CotG is converted into forms of apparent mass ≥ 36 kDa (Fig. S6). We have shown, in previous work, that forms of CotG of apparent mass of 50, 75, and 150 kDa, continue to accumulate following spore release from the mother cell (Zilhao *et al.*, 2005). These species may correspond to phosphorylated forms of CotG, because the co-production of CotG with CotH in *E. coli* (in the absence of CotB) results in the accumulation of forms of the protein with about the same apparent masses (Fig. 4A).

CotH is produced mainly under the control of σ^K^, while the main period of *cotB* and *cotG* expression occurs later, under the control of σ^K^ and GerE (Fig. 1A). Therefore, at least some CotH may already be localized at the spore surface before the onset of the main period of CotB and CotG production. It seems possible that CotB is phosphorylated first, and only then, after spore release from the mother cell, is the full phosphorylation of CotG attained (Zilhao *et al.*, 2005). If phosphorylation of CotG continues following spore release, then it may depend on environmental conditions that control the activity of CotH. Whether or not phosphorylation of CotB-46, CotG and other coat proteins occurs before or after the spore is released from the mother cell, the source of ATP used in the reaction is an unanswered question.

Formation of a normal striated outer coat layer requires CotG and CotH as in spores unable to produce either protein, the outer coat region is expanded, lacks striations and contains disorganized, amorphous material (Fig. 5). The *cotB, cotG*, and *cotH* genes form a functional module that patterns the spore outer coat (Fig. 7C). The expanded outer coat region seen in *cotG* and *cotH* spores bears resemblance to the interspace in spores of the *B. cereus* group, which may also contain coat and exosporium proteins (reviewed by (Stewart, 2015)). It is tempting to suggest that the phosphorylation of ExsB by CotH at Thr residues, versus the phosphorylation of CotG at Ser residues may be part of the reason why the two structural patterns (coat/crust or coat/interspace/exosporium) emerge. We note that ExsB is required for attachment of the exosporium to the coat ((McPherson *et al.*, 2010)), although under certain conditions a reduction or the absence of ExsB may also result in a less robust exosporium located closer to the coat (Aronson *et al.*, 2014).

The resemblance between *cotH* and *cotG* spores is consistent with the lack of accumulation of CotG in the absence of active CotH (Henriques *et al.*, 1998, Saggese *et al.*, 2014). Moreover, it suggests that the main determinant of the striated pattern of the outer coat is CotG and no other CotH-dependent protein (Fig. 7C). Only CotB-46 and CotC seem absent from extracts of *cotG* spores (Sacco *et al.*, 1995, Zilhao *et al.*, 2004); all other outer coat proteins are recruited to the spore surface but lack a scaffold formed by CotG to become organized into a striated pattern ((Henriques *et al.*, 1998); this work). Remarkably, since deletion of the SKR^B^ region has no major impact on the structure of the outer coat, the rationale for polyphosphorylation of CotB-46 remains mysterious. One possibility is that it controls the timing and/or phosphorylation state of CotG (see above).

The role of CotH may be largely confined to patterning of the outer coat, since in *cotB, cotG* or *cotH* mutants the crust layer is still present (Fig. 5). It follows that the normal structure of the outer coat is not a pre-requisite for crust formation (Fig. 7C). In the absence of the crust proteins CotX, CotY and CotZ, however, the outer coat is incomplete. Moreover, the electrondense patches of amorphous material seen in the outer coat region of *cotG* spores form long twirls that project from the spore surface (Fig. 5). Thus, formation of the crust is required for normal assembly of the outer coat, and/or for maintaining the structural integrity and localization of this layer.

The observation that CotZ-GFP formed a complete circle around only about 62% of the WT spores scored, while in the remaining the fusion protein decorated about 2/3 of the circumference of the spore (Fig. 6B), suggests that the crust is a discontinuous, non-uniform structure. CotZ-GFP was found to be enriched at the MCP spore pole when sporulation was induced by resuspension, as we have done here (Imamura *et al.*, 2010), but not when sporulation was induced by resuspension (McKenney *et al.*, 2010, McKenney & Eichenberger, 2012). A non-uniform crust may have functional significance. It calls to mind the assembly of the “bottle cap” in the *B. cereus*/*B. anthracis* group, a specialized spore structure required for germination (Steichen *et al.*, 2007). The cap forms at the MCP pole during the early stages in exosporium assembly before the rest of the exosporium and covers about 1/3 of the spore circumference (Steichen *et al.*, 2007). Strikingly, the *B. subtilis* CotZ and CotY crust proteins are highly similar, and are paralogues of *B. cereus/B. anthracis* CotY and ExsY ((Boydston *et al.*, 2006, Johnson *et al.*, 2006, Redmond *et al.*, 2004, Zhang *et al.*, 1993); reviewed by (Stewart, 2015)). CotY is a cap-specific protein dispensable for assembly of a complete exosporium, whereas ExsY is required for assembly of the non-cap part of the exosporium (Thompson *et al.*, 2012, Thompson & Stewart, 2008, Thompson *et al.*, 2007). Possibly, CotZ and CotY in *B. subtilis* also have specific roles in the assembly of a non-uniform crust.

The widespread occurrence of CotB and CotH orthologues, as well as CotG-like proteins among spore-forming Bacilli (Galperin *et al.*, 2012, Giglio *et al.*, 2011, Saggese *et al.*, 2016) suggests that these proteins could play a role in the organization of the spore surface layers in many species. Differential phosphorylation of homologous proteins may help explaining how a common kit of structural components gives rise to the diverse structural features found at the surface of spores of different spore-forming Firmicutes.

## Material and Methods

### Bacterial strains, media and general techniques

The bacterial strains used in this study are listed in Table S1. Luria-Bertani (LB) medium was routinely used for growth of *E. coli* and *B. subtilis* strains, and sporulation was induced by nutrient exhaustion in liquid Difco sporulation medium (DSM) (Nicholson, 1990). The high fidelity *Phusion* DNA polymerase (Finnzymes) was used in all PCR reactions and the products sequenced to ensure that no unwanted mutations were introduced. Antibiotics were used as described before (Zilhao *et al.*, 2005). All other general methods were as described before (Cutting, 1990, Henriques *et al.*, 1995).

### Plasmids

Details of the construction of all plasmids used in this study can be found in the supplemental material section. The sequence of all primers used is given in Table S2, and all plasmids are listed in Table S3.

### Spore production, purification and spore coat extraction

Mature spores were harvested 24 hours after the onset of sporulation and purified by centrifugation through density gradients of metrizoic acid (Henriques *et al.*, 1995). Proteins were extracted from purified spores using either NaOH or SDS/DTT and fractionated on 12.5% or 15% SDS-PAGE gels as indicated in the figure legends (Henriques *et al.*, 1995). The gels were stained with Coomassie blue R-250 or transferred to nitrocellulose membranes for immunoblotting (described below).

### *B. subtilis* whole cell extracts and immunoblot analysis

Whole cell lysates were prepared from sporulating cultures of *B. subtilis* and resolved by SDS-PAGE (Seyler *et al.*, 1997). σ^A^, CotB, CotG and CotH were immunodetected in spore coat extracts or whole cell lysates using rabbit polyclonal antibodies of established specificity and as previously described (Fujita, 2000, Zilhao *et al.*, 1999, Zilhao *et al.*, 2004). An anti-Phosphoserine antibody was obtained from Milipore and used according to the manufacturer’s guidelines.

### Phosphatase treatment

Proteins were extracted from wild-type purified spores with 0.1M NaOH for 15 min at 4 °C. After centrifugation at 12,000*g*, for 10 min at 4 °C the supernatant containing the coat proteins was neutralized with 0.1M HCl. The lysate was then subject to treatment with alkaline phosphatase (FastAP, from Fermentas); 2 enzyme units were added to 50 µl of neutralized extract and the mixture incubated at 37°C for 20 min, prior to addition of SDS-PAGE loading dye to stop the reaction.

### Overproduction of CotB-His_6_, CotG-His_6_ and CotH

The various recombinant proteins were overexpressed in *E. coli* BL21(DE3) by a modified auto-induction method (Fernandes *et al.*, 2015). After the induction period, cells were collected by centrifugation (at 12,000*g*, for 10 min at 4 °C) and resuspended in one tenth of the culture volume of buffer W (100 mM Tris-HCl, pH 8.0 and 150 mM NaCl). The cells were then lysed by passage through a French Press cell at 19,000 lb/in^2^ as described previously (Henriques *et al.*, 1995). Samples of the whole cell lysates were electrophoretically resolved ony 12% SDS–PAGE. Staining of SDS-PAGE gels with Pro-Q^®^ Diamond and SYPRO^®^ Ruby staining was as described by the manufacturer (Invitrogen). The PeppermintStick phosphoprotein molecular weight standard was also used as described by the manufacturer (Invitrogen); in this marker, ovalbumin (45 kDa) and β-casein (23.6 kDa) are phosphorylated. A FLA-5100 fluorescence scanner (from Fuji) was used for imaging of the stained gels.

### Purification of CotB, CotH and CotH^D228Q^

CotB-His_6_ and CotH-*Strep-tag* proteins were overexpressed in *E. coli* BL21(DE3) as described above (Fernandes *et al.*, 2015). After the induction period, cells were collected by centrifugation (at 12,000*g*, for 10 min at 4 °C). Cells with CotB-His_6_ were resuspended in one tenth of Start buffer (10 mM imidazole, 20 mM phosphate, 0.5 M NaCl), containing 1 M phenylmethanesulfonyl fluoride (PMSF). Lysates were prepared by a passage through a French presys (19000 psi) and centrifuged (at 12,000*g*, for 10 min at 4 °C). Since most of CotB-His_6_ is present in the insoluble fraction, Start buffer containing urea (8 M) was added to the debris to increase the solubility of CotB-His_6_. The sample was stirred for 45 min and centrifuged (at 12,000*g*, for 10 min). CotB-His_6_ was purified by Ni^2+^-NTA affinity chromatography (Qiagen). The Ni^2+^-NTA-affinity purified protein was analyzed by 12.5% SDS-PAGE. Fractions containing CotB-His_6_ were pooled and dialyzed against Start buffer without ureia. Cells with CotH-*Strep-tag* or CotH^D228Q^ –*Strep-tag* were resuspended in one tenth of buffer W (see above), containing 1 M PMSF. Lysates were prepared by a passage through a French press (19000 psi) and centrifuged (at 12,000*g*, for 10 min at 4 °C). CotH-*Strep-tag* or CotH^D228Q^– *Strep-tag* were purified using Strep-Tactin Sepharose (IBA). The affinity purified protein was analyzed by 12.5% SDS-PAGE.

### Kinase activity assays

Purified CotH (0.5-1μM) and/or CotB (2μM) were incubated with 0.1 mM of unlabelled ATP and 1 □Ci of [γ -^32^P]ATP in kinase buffer (50 mM Tris-HCl pH 7.5, 50 mM KCl, 10 mM MgCl_2_, 10 mM MnCl_2_ and 0.5 mM TCEP) at 37°C for 5, 10 and 30 min. For kinase inhibition, staurosporine (Sigma) was added at concentrations of 7, 70 or 700 μM. The reactions were stopped by addition of SDS-PAGE loading buffer and the proteins resolved by SDS-PAGE. The gels were dried, exposed to a Phospho Screen and imaged using a Storm phosphorimager (GE Healthcare).

### Germination efficiency

Purified spores were heat activated as previously described (Cutting, 1990) and diluted in 10 mM Tris-HCl pH 8.0 buffer containing 1 mM glucose, 1 mM fructose, and 10 mM KCl. After 15 min at 37 °C, L-asparagine was added to a final concentration of 10 mM, and the optical density of the suspension at 580 nm ion was measured at 10 min intervals until a constant reading was reached.

### Transmission Electron Microscopy

For thin sectioning transmission electron microscopy (TEM) analysis, *B. subtilis* spores were purified by density gradient centrifugation as described above. Samples were processed for TEM essentially as described previously (Henriques *et al.*, 1998) and imaged on a Hitachi H-7650 Microscope equipped with an AMT digital camera operated at 120 keV.

### Mass spectrometry

Cultures of *B. subtilis* cells were induced to sporulate by resuspension in Sterlini–Mandelstam (SM) medium (Please cite PMID: 2104613) and samples were taken every hour for five hours. Then, the cells were sedimented at 4°C and 8,000xg for 10 minutes. The cell pellets were resuspended in 20 ml of lysis/binding buffer (8M Urea, 20mM Tris-HCl pH 8.0, 150mM NaCl, 1mM PMSF, 1mM β–glycerol phosphate, 1mM sodium orthovanadate, 2.5mM sodium pyrophosphate and 10mM sodium fluoride) and disrupted using a cell-disruptor (Constant Systems Limited). The lysate was centrifuged for 30 minutes at 15,000xg to precipitate the cell debris. Then, free cysteines were alkylated with 5mM Iodoacetamide and incubated for 30 minutes in the dark. Then, the urea concentration was decreased to below 1M by the addition of 50 mM Tris/HCl pH 8 and 150 mM NaCl and proteins were subsequently digested with trypsin (1:20, Promega) for 12 h. Then, phospho-peptide enrichment was performed using Titansphere TiO2 beads (GL Science) according to a previously described protocol (please cite PMID:22517742). Briefly, the peptide mix was acidified using 1 % TFA and solution was cleared by centrifugation for 5 minutes at 15,000xg at room temperature. The supernatant was purified using a C18 Sep-Pak columns (Waters) and eluted with 50 % acetonitrile and 0.1 %TFA. Samples were lyophilized and resuspendend in 73 % Acetonitrile, 25 % lactic acid and 2 % TFA, mixed with 30 µg beads (GLC Science) and incubated at room temperature for 30 minutes. Beads were loaded onto a C18 spin column (Nest group) and washed three times with 80 % acetonitrile and 2 % TFA. Peptides were eluted with 50 µl 5% NH_4_OH followed by 50 µl 50 % acetonitrile.

Enriched peptides were separated and analyzed by LC-MS/MS using an Easy-nLCII HPLC system (Thermo Fisher Scientific) coupled directly to an LTQ Orbitrap Velos mass spectrometer (Thermo Fisher Scientific) at the Harvard Mass Spectrometry and Proteomics Resource Laboratory, FAS Center for Systems Biology, Northwest Bldg. Room B247, 52 Oxford St, Cambridge, MA.

## Supporting information

## Acknowledgements

We acknowledge A.L. Sousa and E.M. Tranfield from the Electron Microscopy Facility at the Instituto Gulbenkian de Ciência for sample processing and technical expertise; we thank Ana Henriques for art work. This work was financially supported by Project LISBOA-01-0145-FEDER-007660 (“Microbiologia Molecular, Estrutural e Celular”) funded by FEDER funds through COMPETE2020 – “Programa Operacional Competitividade e Internacionalização” (POCI), by National Institutes of Health Grant NIH GM18568 to R. Losick, and through FCT (“Fundação para a Ciência e a Tecnologia”) grants POCI/BIA-BCM/60855/2004 and POCTI/BCI/48647/2002 to A.O.H. and programme IF (IF/00268/2013/CP1173/CT0006) to M.S.

## References

Abhyankar, W., A.H. Hossain, A. Djajasaputra, P. Permpoonpattana, A. Ter Beek, H.L. Dekker, S.M. Cutting, S. Brul, L.J. de Koning & C.G. de Koster, (2013) In pursuit of protein targets: proteomic characterization of bacterial spore outer layers. J Proteome Res 12: 4507–4521.

Abhyankar, W., S. Stelder, L. de Koning, C. de Koster & S. Brul, (2017) ‘Omics’ for microbial food stability: Proteomics for the development of predictive models for bacterial spore stress survival and outgrowth. Int J Food Microbiol 240: 11–18.

Aronson, A., B. Goodman & Z. Smith, (2014) The regulated synthesis of a Bacillus anthracis spore coat protein that affects spore surface properties. J Appl Microbiol 116: 1241–1249.

Baccigalupi, L., G. Castaldo, G. Cangiano, R. Isticato, R. Marasco, M. De Felice & E. Ricca, (2004) GerE-independent expression of cotH leads to CotC accumulation in the mother cell compartment during Bacillus subtilis sporulation. Microbiology 150: 3441– 3449.

Ball, D.A., R. Taylor, S.J. Todd, C. Redmond, E. Couture-Tosi, P. Sylvestre, A. Moir & P.A. Bullough, (2008) Structure of the exosporium and sublayers of spores of the Bacillus cereus family revealed by electron crystallography. Mol Microbiol 68: 947–958.

Boydston, J.A., L. Yue, J.F. Kearney & C.L. Turnbough, Jr., (2006) The ExsY protein is required for complete formation of the exosporium of Bacillus anthracis. J Bacteriol 188: 7440–7448.

Costa, T., M. Serrano, L. Steil, U. Volker, C.P. Moran, Jr. & A.O. Henriques, (2007) The timing of cotE expression affects Bacillus subtilis spore coat morphology but not lysozyme resistance. J Bacteriol 189: 2401–2410.

Cutting, S.a.H.P.B., (1990) Genetic Analysis. John Wiley and sons, Lta.

Donovan, W., L.B. Zheng, K. Sandman & R. Losick, (1987) Genes encoding spore coat polypeptides from Bacillus subtilis. J Mol Biol 196: 1–10.

Driks, A. & P. Eichenberger, (2016) The Spore Coat. Microbiol Spectr 4.

Fernandes, C.G., D. Placido, D. Lousa, J.A. Brito, A. Isidro, C.M. Soares, J. Pohl, M.A. Carrondo, M. Archer & A.O. Henriques, (2015) Structural and Functional Characterization of an Ancient Bacterial Transglutaminase Sheds Light on the Minimal Requirements for Protein Cross-Linking. Biochemistry 54: 5723–5734.

Fujita, M., (2000) Temporal and selective association of multiple sigma factors with RNA polymerase during sporulation in Bacillus subtilis. Genes Cells 5: 79–88.

Galperin, M.Y., S.L. Mekhedov, P. Puigbo, S. Smirnov, Y.I. Wolf & D.J. Rigden, (2012) Genomic determinants of sporulation in Bacilli and Clostridia: towards the minimal set of sporulation-specific genes. Environ Microbiol 14: 2870–2890.

Giglio, R., R. Fani, R. Isticato, M. De Felice, E. Ricca & L. Baccigalupi, (2011) Organization and evolution of the cotG and cotH genes of Bacillus subtilis. J Bacteriol 193: 6664–6673.

Giorno, R., J. Bozue, C. Cote, T. Wenzel, K.S. Moody, M. Mallozzi, M. Ryan, R. Wang, R. Zielke, J.R. Maddock, A. Friedlander, S. Welkos & A. Driks, (2007) Morphogenesis of the Bacillus anthracis spore. J Bacteriol 189: 691–705.

Henriques, A.O., B.W. Beall, K. Roland & C.P. Moran, Jr., (1995) Characterization of cotJ, a sigma E-controlled operon affecting the polypeptide composition of the coat of Bacillus subtilis spores. J Bacteriol 177: 3394–3406.

Henriques, A.O., L.R. Melsen & C.P. Moran, Jr., (1998) Involvement of superoxide dismutase in spore coat assembly in Bacillus subtilis. J Bacteriol 180: 2285–2291.

Henriques, A.O. & C.P. Moran, Jr., (2007) Structure, assembly, and function of the spore surface layers. Annu Rev Microbiol 61: 555–588.

Imamura, D., R. Kuwana, H. Takamatsu & K. Watabe, (2010) Localization of proteins to different layers and regions of Bacillus subtilis spore coats. J Bacteriol 192: 518–524.

Isticato, R., G. Esposito, R. Zilhao, S. Nolasco, G. Cangiano, M. De Felice, A.O. Henriques & E. Ricca, (2004) Assembly of multiple CotC forms into the Bacillus subtilis spore coat. J Bacteriol 186: 1129–1135.

Isticato, R., A. Pelosi, R. Zilhao, L. Baccigalupi, A.O. Henriques, M. De Felice & E. Ricca, (2008) CotC-CotU heterodimerization during assembly of the Bacillus subtilis spore coat. J Bacteriol 190: 1267–1275.

Isticato, R., T. Sirec, R. Giglio, L. Baccigalupi, G. Rusciano, G. Pesce, G. Zito, A. Sasso, M. De Felice & E. Ricca, (2013) Flexibility of the programme of spore coat formation in Bacillus subtilis: bypass of CotE requirement by over-production of CotH. PLoS One 8: e74949.

Isticato, R., T. Sirec, S. Vecchione, A. Crispino, A. Saggese, L. Baccigalupi, E. Notomista, A. Driks & E. Ricca, (2015) The Direct Interaction between Two Morphogenetic Proteins Is Essential for Spore Coat Formation in Bacillus subtilis. PLoS One 10: e0141040.

Jiang, S., Q. Wan, D. Krajcikova, J. Tang, S.B. Tzokov, I. Barak & P.A. Bullough, (2015) Diverse supramolecular structures formed by self-assembling proteins of the Bacillus subtilis spore coat. Mol Microbiol 97: 347–359.

Johnson, M.J., S.J. Todd, D.A. Ball, A.M. Shepherd, P. Sylvestre & A. Moir, (2006) ExsY and CotY are required for the correct assembly of the exosporium and spore coat of Bacillus cereus. J Bacteriol 188: 7905–7913.

Kailas, L., C. Terry, N. Abbott, R. Taylor, N. Mullin, S.B. Tzokov, S.J. Todd, B.A. Wallace, J.K. Hobbs, A. Moir & P.A. Bullough, (2011) Surface architecture of endospores of the Bacillus cereus/anthracis/thuringiensis family at the subnanometer scale. Proc Natl Acad Sci U S A 108: 16014–16019.

Kim, H., M. Hahn, P. Grabowski, D.C. McPherson, M.M. Otte, R. Wang, C.C. Ferguson, P. Eichenberger & A. Driks, (2006) The Bacillus subtilis spore coat protein interaction network. Mol Microbiol 59: 487–502.

Liu, Z. & Y. Huang, (2014) Advantages of proteins being disordered. Protein Sci 23: 539–550.

McKenney, P.T., A. Driks & P. Eichenberger, (2013) The Bacillus subtilis endospore: assembly and functions of the multilayered coat. Nat Rev Microbiol 11: 33–44.

McKenney, P.T., A. Driks, H.A. Eskandarian, P. Grabowski, J. Guberman, K.H. Wang, Z. Gitai & P. Eichenberger, (2010) A distance-weighted interaction map reveals a previously uncharacterized layer of the Bacillus subtilis spore coat. Curr Biol 20: 934– 938.

McKenney, P.T. & P. Eichenberger, (2012) Dynamics of spore coat morphogenesis in Bacillus subtilis. Mol Microbiol 83: 245–260.

McPherson, S.A., M. Li, J.F. Kearney & C.L. Turnbough, Jr., (2010) ExsB, an unusually highly phosphorylated protein required for the stable attachment of the exosporium of Bacillus anthracis. Mol Microbiol 76: 1527–1538.

Naclerio, G., L. Baccigalupi, R. Zilhao, M. De Felice & E. Ricca, (1996) Bacillus subtilis spore coat assembly requires cotH gene expression. J Bacteriol 178: 4375–4380.

Nguyen, K.B., A. Sreelatha, E.S. Durrant, J. Lopez-Garrido, A. Muszewska, M. Dudkiewicz, M. Grynberg, S. Yee, K. Pogliano, D.R. Tomchick, K. Pawlowski, J.E. Dixon & V.S. Tagliabracci, (2016) Phosphorylation of spore coat proteins by a family of atypical protein kinases. Proc Natl Acad Sci U S A 113: E3482–3491.

Nicholson, W.L.a.S. P., (1990) Sporulation, Germination and Outgrowth. In: Molecular Biology Methods for Bacillus. H.C.R.a.C. S.M. (ed). Chichester: John Wiley & Sons Ltd, pp. 391–450.

Ramamurthi, K.S. & R. Losick, (2008) ATP-driven self-assembly of a morphogenetic protein in Bacillus subtilis. Mol Cell 31: 406–414.

Redmond, C., L.W. Baillie, S. Hibbs, A.J. Moir & A. Moir, (2004) Identification of proteins in the exosporium of Bacillus anthracis. Microbiology 150: 355–363.

Romero, Obradovic & K. Dunker, (1997) Sequence Data Analysis for Long Disordered Regions Prediction in the Calcineurin Family. Genome Inform Ser Workshop Genome Inform 8: 110–124.

Rosenberg, A., B. Soufi, V. Ravikumar, N.C. Soares, K. Krug, Y. Smith, B. Macek & S. Ben-Yehuda, (2015) Phosphoproteome dynamics mediate revival of bacterial spores. BMC Biol 13: 76.

Ruegg, U.T. & G.M. Burgess, (1989) Staurosporine, K-252 and UCN-01: potent but nonspecific inhibitors of protein kinases. Trends Pharmacol Sci 10: 218–220.

Sacco, M., E. Ricca, R. Losick & S. Cutting, (1995) An additional GerE-controlled gene encoding an abundant spore coat protein from Bacillus subtilis. J Bacteriol 177: 372– 377.

Saggese, A., R. Isticato, G. Cangiano, E. Ricca & L. Baccigalupi, (2016) CotG-Like Modular Proteins Are Common among Spore-Forming Bacilli. J Bacteriol 198: 1513– 1520.

Saggese, A., V. Scamardella, T. Sirec, G. Cangiano, R. Isticato, F. Pane, A. Amoresano, E. Ricca & L. Baccigalupi, (2014) Antagonistic role of CotG and CotH on spore germination and coat formation in Bacillus subtilis. PLoS One 9: e104900.

Setlow, P., (2014) Germination of spores of Bacillus species: what we know and do not know. J Bacteriol 196: 1297–1305.

Setlow, P., S. Wang & Y.Q. Li, (2017) Germination of Spores of the Orders Bacillales and Clostridiales. Annu Rev Microbiol 71: 459–477.

Seyler, R.W., Jr., A.O. Henriques, A.J. Ozin & C.P. Moran, Jr., (1997) Assembly and interactions of cotJ-encoded proteins, constituents of the inner layers of the Bacillus subtilis spore coat. Mol Microbiol 25: 955–966.

Shah, I.M., M.H. Laaberki, D.L. Popham & J. Dworkin, (2008) A eukaryotic-like Ser/Thr kinase signals bacteria to exit dormancy in response to peptidoglycan fragments. Cell 135: 486–496.

Steichen, C.T., J.F. Kearney & C.L. Turnbough, Jr., (2007) Non-uniform assembly of the Bacillus anthracis exosporium and a bottle cap model for spore germination and outgrowth. Mol Microbiol 64: 359–367.

Stewart, G.C., (2015) The Exosporium Layer of Bacterial Spores: a Connection to the Environment and the Infected Host. Microbiol Mol Biol Rev 79: 437–457.

Thompson, B.M., B.C. Hoelscher, A. Driks & G.C. Stewart, (2012) Assembly of the BclB glycoprotein into the exosporium and evidence for its role in the formation of the exosporium ‘cap’ structure in Bacillus anthracis. Mol Microbiol 86: 1073–1084.

Thompson, B.M. & G.C. Stewart, (2008) Targeting of the BclA and BclB proteins to the Bacillus anthracis spore surface. Mol Microbiol 70: 421–434.

Thompson, B.M., L.N. Waller, K.F. Fox, A. Fox & G.C. Stewart, (2007) The BclB glycoprotein of Bacillus anthracis is involved in exosporium integrity. J Bacteriol 189: 6704–6713.

Todd, S.J., A.J. Moir, M.J. Johnson & A. Moir, (2003) Genes of Bacillus cereus and Bacillus anthracis encoding proteins of the exosporium. J Bacteriol 185: 3373–3378.

Xiao, J., V.S. Tagliabracci, J. Wen, S.A. Kim & J.E. Dixon, (2013) Crystal structure of the Golgi casein kinase. Proc Natl Acad Sci U S A 110: 10574–10579.

Zhang, J., P.C. Fitz-James & A.I. Aronson, (1993) Cloning and characterization of a cluster of genes encoding polypeptides present in the insoluble fraction of the spore coat of Bacillus subtilis. J Bacteriol 175: 3757–3766.

Zilhao, R., R. Isticato, L.O. Martins, L. Steil, U. Volker, E. Ricca, C.P. Moran, Jr. & A.O. Henriques, (2005) Assembly and function of a spore coat-associated transglutaminase of Bacillus subtilis. J Bacteriol 187: 7753–7764.

Zilhao, R., G. Naclerio, A.O. Henriques, L. Baccigalupi, C.P. Moran, Jr. & E. Ricca, (1999) Assembly requirements and role of CotH during spore coat formation in Bacillus subtilis. J Bacteriol 181: 2631–2633.

Zilhao, R., M. Serrano, R. Isticato, E. Ricca, C.P. Moran, Jr. & A.O. Henriques, (2004) Interactions among CotB, CotG, and CotH during assembly of the Bacillus subtilis spore coat. J Bacteriol 186: 1110–1119.

