## Supplementary material for "A protein phosphorylation module patterns the *Bacillus subtilis* spore outer coat"

#### Supplemental Material

#### Supplemental Material and Methods

**Construction of a *cotH*$^{D228Q}$ allele.** PCR was used to fuse the promoter region that regulates expression of *cotH* to the coding region, with the consequent elimination of the *cotG* gene present within the 5´-unstralated region of the cotH mRNA (Giglio *et al.*, 2011). First, the promoter downstream of *cotG* (449 bp) was amplified with primers cotH95D and cotGcotHR, and the promoter upstream of *cotH* was amplified together with *cotH* with primers cotGcotHD and cotH 2140R (1329 bp). Both fragments were mixed and the resulting fragment of 1778 bp was amplified using primer cotH95D and cotH2104R. The fragment was digested with SalI and HindIII and cloned between the same sites of pCD6, to yield pCD7. pCD6 is a derivative of pMLK83 (Karow & Piggot, 1995), in which the *gusA* gene was removed by digestion with EcoRI. We then used primers *cotH* D228Q_D and D228Q_R to replace the aspartate codon 228 to a glutamine codon in the activation loop of CotH, to produce pCD8.

**Construction of *E. coli* strains for the overproduction of Cot proteins.** A 601 bp PCR fragment encoding *cotG* was generated with primers G/5/Nco and *cotG*-R/NotI. The resulting fragment was cleaved with *NcoI* and *NotI* and inserted between the same sites of pACYCDuet-1 (Novagen) to yield pAP39. Primers *cotH*-R/*XhoI* and *cotH*-F/*NdeI* were used to PCR amplify *cotH* coding region. The resulting 1,122 bp fragment was cleaved with *XhoI* and *NdeI* and cloned between the same sites of pACYCDuet-1 to yield pCD10. pCD11 was generated by cloning the cleaved (*XhoI*-*NdeI*) *cotH* fragment between the same sites of pAP39. Primers *cotH* D228Q_D and D228Q_R were used with pCD10 and pCD11 as the templates, to replace the aspartate codon 228 to a glutamine codon in the activation loop of CotH, to produce pNP5 and pNP6, respectively.

**Yeast two-hybrid analysis.** The complete coding regions of *cotB* and *cotG* were PCR amplified with primers *cotB*-5/Nco and *cotB*-OMO/3 and primers *cotG*-5/*Nco* and *cotG*-*Bam*/3 (Table S2). The 5´ (coding for residues 1-213) and 3´-end regions (encoding amino acids 195-380) of *cotB* were amplified separately using primers *cotB*-5/*Nco* and *cotB*-OMO/6 and *cotB*-*Nco*/ID and *cotB*-OMO/3, respectively (Table S2). The entire *cotB*, as well as its N- and C-terminal regions, and *cotG* PCR products were digested by *Nco*I and *Bam*HI and inserted between the *Nco*I and *Bam*HI sites of both pAS2-1 and pACT2 (Clontech), to create in frame fusions to the GAL4 DNA binding (DNA-BD) or activation domains (AD). The *cotH* coding sequence was obtained by PCR using two different sets of primers *cotH*/5/Nde and *cotH*/3/*Bam* and *cotH*/5/*Bam* and *cotH*/3/*Xho* (Table S2). The *cotH* PCR products were digested with *Nde*I and *Bam*HI and inserted between the same sites of pAS2-1, or with *Bam*HI and *Xho*I and fused to the GAL4 AD in pACT2. Yeast strains Y187 (MAT$\alpha$, *ura*3-52, *his*3-200, *ade*2-101, *trp*1-901, *leu*2-3, 112, *gal*4$\Delta$, *met-, gal*80$\Delta$, URA3::GAL1$_{UAS}$-GAL1$_{TATA}$-*lacZ*) and Y190 (MATa, *ura*3-52, *his*3-200, *ade*2-101, *lys*2-801, *trp*1-901, *leu*2-3, 112, *gal*4$\Delta$, *gal*80$\Delta$, cyh$^r$2, LYS2::URA::GAL1$_{UAS}$-HIS3$_{TATA}$-HIS3, URA3::GAL1$_{UAS}$-GAL1$_{TATA}$-*lacZ*) (Matchmaker Two-hybrid system, Clontech) were transformed independently with the pAS2-1 and pACT-2 vectors and/or each of these constructs respectively. The resulting clones were used in pairwise matings selecting for LEU$^+$ and TRP$^+$. Colony lift assays assays for detection of β-galactosidase activity were essentially as previously described (Ozin *et al.*, 2000).

### Supplemental Results and Discussion

#### Phosphosites in CotG

Saggese and co-authors found pSer at position 15 of CotG, outside the SKR$^{G}$ region, as well as pSer at position 39 and pThr at position 147 within the SKR$^{G}$ region (Saggese *et al.*, 2016) (Fig. 1C). In addition, 9 tripeptides within the SKR$^{G}$ region contained a phosphate moiety although the phosphorylated residue could not be unambiguously identified (Saggese *et al.*, 2014) (Fig. 1C). We confirmed the identification of pSer39, we unambiguously identified pSer at 4 of these tripeptides, and we additionally found 3 other pSer in the SKR$^{G}$ region, but have not detected pSer15 (Fig. 1C). Moreover, purified CotH phosphorylated a peptide derived from the SKR$^{G}$ region (residues 88 to 100, encompassing part of R8, R9 and part of R10) at Ser88 or Ser90 (according to the numbering of Fig. 1C, for the full-length protein) (Nguyen *et al.*, 2016). Phosphorylation of Ser88 was not detected in our study. Thus, CotG may be phosphorylated at a minimum of 15-16 positions: at Ser15, and at 14-15 residues within the SKR$^{G}$ region, 13-14 of which are pSer (verified or most likely) and 1 is pThr147 ((Nguyen *et al.*, 2016, Saggese *et al.*, 2014); this work) (Fig. 1C). That most phosphorylation sites in CotB and CotG are in the SKR$^{B}$ and SKR$^{G}$ regions is in line with the finding that protein phosphorylation predominates in regions of intrinsic sequence disorder (Collins *et al.*, 2008).

#### Spores of a *cotH*$^{D228Q}$ mutant are impaired in germination

We monitored the germination response of WT spores and those of the *cotH* insertional mutant, in parallel with spores of the insertional mutant bearing either the WT or *cotH*$^{D228Q}$ alleles at *amyE*. Spores formed by the *cotH* insertional or *cotHD288Q* point mutants were strongly impaired in germination in response to AGFK (Fig. S2B). Interestingly, the presence of the inactive kinase, in *cotHD288Q* spores, caused a greater impairment in AGFK-triggered germination than the absence of the kinase, in the deletion mutant (Fig. S2B). Possibly, the absence of the CotH protein allows another kinase to partially replace it, whereas the presence of $\text{CotH}^{D228Q}$ excludes this putative second kinase. We note that the *yisJ* gene, located just upstream of the $\sigma^K$-controlled *gerP* locus, required for normal spore germination (Behravan *et al.*, 2000, Butzin *et al.*, 2012), codes for a kinase with significant sequence similarity to CotH (not shown). *yisJ* may also be involved in coat assembly and proper spore germination (our unpublished results). In any event, and in agreement with previous results (Nguyen *et al.*, 2016), the activity of CotH is required for proper spore germination.

**Phosphorylation of CotB-46 and CotG at the spore surface**

Transcription of *cotH* occurs mainly under the control of $\sigma^K$, whereas most of the *cotB* and *cotG* transcription is $\sigma^K$ and GerE-dependent and thus occurs later (Eichenberger *et al.*, 2004, Eichenberger *et al.*, 2003, Sacco *et al.*, 1995, Steil *et al.*, 2005). This raises the possibility that CotH is assembled before CotG and CotB are produced, and thus that the phosphorylation reactions involving these proteins occur at the spore surface. To test this idea, sporulating cells of a wild-type strain were harvested at various times during sporulation, lysed, and mother cell and forespore fractions prepared by centrifugation (Zilhao *et al.*, 2004). The presence of CotB-46, CotB-66 and CotG in either fraction was then monitored by immunoblotting (Zilhao *et al.*, 2004). CotB-66 was detected at hour 6 of sporulation in the forespore fraction of a WT strain (Fig. S6). From this time of sporulation onwards, only CotB-66 was detected suggesting that CotB-46 can be phosphorylated at the spore surface (Fig. S6). Neither form of CotB was detected in the mother cell fraction at any time point tested, even though the amount of total protein analysed (30 µg) was larger than for the forespore fraction (10 µg). Thus, both CotB-46 and CotB-66 appear enriched in the forespore fraction. Only CotB-46 was detected in the forespore fraction of a *cotG* mutant, in line with the observation that *cotG* is not required for the assembly of CotB-46 but promotes formation of CotB-66 ((Zilhao *et al.*, 2004); this work). CotG-24 may be rapidly converted into CotG-36, which was the first form of the protein detected, at hour 7, and only in the forespore fraction (Fig. S6). With time, forms of CotG around 50 kDa, 75 kDa and 100 kDa become visible in the spore fraction. Phosphorylation of CotB-46 and of CotG in the mother cell may have not been detected with our temporal resolution. In any case, phosphorylation of both CotB-46 and CotG can occur at the spore surface.

### References


Behravan, J., H. Chirakkal, A. Masson & A. Moir, (2000) Mutations in the gerP locus of Bacillus subtilis and Bacillus cereus affect access of germinants to their targets in spores. *J Bacteriol* **182**: 1987-1994.

Butzin, X.Y., A.J. Troiano, W.H. Coleman, K.K. Griffiths, C.J. Doona, F.E. Feeherry, G. Wang, Y.Q. Li & P. Setlow, (2012) Analysis of the effects of a gerP mutation on the germination of spores of Bacillus subtilis. *J Bacteriol* **194**: 5749-5758.

Collins, M.O., L. Yu, I. Campuzano, S.G. Grant & J.S. Choudhary, (2008) Phosphoproteomic analysis of the mouse brain cytosol reveals a predominance of protein phosphorylation in regions of intrinsic sequence disorder. *Mol Cell Proteomics* **7**: 1331-1348.

Eichenberger, P., M. Fujita, S.T. Jensen, E.M. Conlon, D.Z. Rudner, S.T. Wang, C. Ferguson, K. Haga, T. Sato, J.S. Liu & R. Losick, (2004) The program of gene transcription for a single differentiating cell type during sporulation in Bacillus subtilis. *PLoS Biol* **2**: e328.

Eichenberger, P., S.T. Jensen, E.M. Conlon, C. van Ooij, J. Silvaggi, J.E. Gonzalez-Pastor, M. Fujita, S. Ben-Yehuda, P. Stragier, J.S. Liu & R. Losick, (2003) The sigmaE regulon and the identification of additional sporulation genes in Bacillus subtilis. *J Mol Biol* **327**: 945-972.

Giglio, R., R. Fani, R. Isticato, M. De Felice, E. Ricca & L. Baccigalupi, (2011) Organization and evolution of the cotG and cotH genes of Bacillus subtilis. *J Bacteriol* **193**: 6664-6673.

Karow, M.L. & P.J. Piggot, (1995) Construction of gusA transcriptional fusion vectors for Bacillus subtilis and their utilization for studies of spore formation. *Gene* **163**: 69-74.

Nguyen, K.B., A. Sreelatha, E.S. Durrant, J. Lopez-Garrido, A. Muszewska, M. Dudkiewicz, M. Grynberg, S. Yee, K. Pogliano, D.R. Tomchick, K. Pawlowski, J.E. Dixon & V.S. Tagliabracci, (2016) Phosphorylation of spore coat proteins by a family of atypical protein kinases. *Proc Natl Acad Sci U S A* **113**: E3482-3491.

Ozin, A.J., A.O. Henriques, H. Yi & C.P. Moran, Jr., (2000) Morphogenetic proteins SpoVID and SafA form a complex during assembly of the Bacillus subtilis spore coat. *J Bacteriol* **182**: 1828-1833.

Sacco, M., E. Ricca, R. Losick & S. Cutting, (1995) An additional GerE-controlled gene encoding an abundant spore coat protein from Bacillus subtilis. *J Bacteriol* **177**: 372-377.

Saggese, A., R. Isticato, G. Cangiano, E. Ricca & L. Baccigalupi, (2016) CotG-Like Modular Proteins Are Common among Spore-Forming Bacilli. *J Bacteriol* **198**: 1513-1520.

Saggese, A., V. Scamardella, T. Sirec, G. Cangiano, R. Isticato, F. Pane, A. Amoresano, E. Ricca & L. Baccigalupi, (2014) Antagonistic role of CotG and CotH on spore germination and coat formation in Bacillus subtilis. *PLoS One* **9**: e104900.

Steil, L., M. Serrano, A.O. Henriques & U. Volker, (2005) Genome-wide analysis of temporally regulated and compartment-specific gene expression in sporulating cells of Bacillus subtilis. *Microbiology* **151**: 399-420.

Zilhao, R., M. Serrano, R. Isticato, E. Ricca, C.P. Moran, Jr. & A.O. Henriques, (2004) Interactions among CotB, CotG, and CotH during assembly of the Bacillus subtilis spore coat. *J Bacteriol* **186**: 1110-1119.

### Supplemental figure legends

**Figure S1 – Disorder prediction and distribution of serine residues in CotB and CotG. A**: organization of CotB and CotG, with relevant regions represented in color: red and blue for the NTD domain and repeat domains of CotB, and yellow for the repeat region of CotG. Below the representation of the proteins, is a prediction of order/disorder along the primary structures of CotB and CotG. The plots were generated with the PONDR program (www.pondr.org). Regions with scores above 0.5 are most likely disordered. Two long stretches (black bars) in CotB and CotG, that are most likely disordered, coincide with their repeat regions. **B**: prediction of the propensity of serine, threonine, and tyrosine residues to be phosphorylated along the primary structures of CotB and CotG. The plots were generated with the DIPHOS program (www.diphos.org). CotB has 50 serine residues showing a DIPHOS score of 1.0, all of which are located within the SKR$^{B}$ region; 5 other serine residues at the C-terminus of the repeat region show a lower probability of phosphorylation. CotG has 35 serine residues with a DIPHOS score close to one, all of which located inside the SKR$^{G}$ region; three additional serine residues upstream of the repeat region, and on downstream of this region show a lower DIPHOS score.

**Figure S2 – Assembly of CotH$^{D228Q}$. A:** Spores produced by a wild type (WT) strain, by *cotB*, *cotG*, *cotH* mutants and by a *cotH* mutant with either *cotH*$^{wt}$ or *cotH*$^{D228Q}$ at *amyE*, were purified. The coat proteins were extracted by SDS-DTT treatment, analysed by SDS-PAGE and Coomassie staining. **B**: *cotH*$^{D228Q}$ spores show impaired germination. Spores produced by a wild type (WT) strain, by a *cotH* mutant and by a Δ*cotH* mutant with either *cotHwt* or *cotHD228Q* at *amyE*, were purified, and the kinetics of germination was examined. Germination was induced by L-aspargine in GFK, and monitored by the decrease in OD$_{600}$ of the spore suspension. The efficiency of germination is defined as the ratio between the optical density of the culture at a given time after exposure to the germinant mixture and the original optical density of the suspension.

**Figure S3 – Pro-Q Diamond staining to assess phosphorylation of CotB and CotG in *E. coli*. A** and **B** correspond to panels A and B of Fig. 3, respectively, but the gels are shown stained with Sypro Ruby (top) or Pro-Q Diamond (bottom). The Peppermint phosphoprotein marker (Pe) was used, in which bands *b* (ovalbumin) and *c* (β-casein) are phosphorylated, while band *a* (BSA) is not. The bottom part of the panel shows a black and white inverted image of the marker lanes stained with Sypro Ruby (S) and Pro-Q Diamond (Q), and the quantification of the Q/S signal for the *a* and *b* proteins as well as for CotB-46 (B), CotB-66 (B+G+H), and the intermediate form of CotB produced in the presence of CotB and CotH alone (B+H) (see also Figure 3). The position of molecular weight markers is shown on the left side of the panels.

**Figure S4 – CotH-Strep-tag has kinase activity. A:** organization of CotH with the kinase domain represented in brown and the Strep-*tag* in yellow. **B**: *E. coli* strains producing the indicated proteins from T7*lac* promoters were grown in autoinduction medium and whole cell extracts prepared. Proteins in the extracts were resolved by SDS-PAGE and the gels stained with Coomassie Blue. The position of CotH (green arrow), CotB-66 (blue arrow) and CotB-46 (red arrow) are indicated. A brown arrow shows the position of a form of CotB with an apparent mass between that of CotB-46 and CotB-66. The position of MW marker proteins (in kDa) is shown on the left side of the pannel.

**Figure S5 – Autophosphorylation and transphophorylation activity of CotH is insensitive to staurosporine. A:** Purified CotH and CotH$^{D228Q}$, at the indicated concentrations, incubated in the presence of [γ-32P]ATP for 0 or 30 minutes. In the right panels staurosporine (STP) was added at the indicated concentration to the phosphorylation reaction. The reactions were stopped by adding loading buffer and were run in a SDS-PAGE, phosphorylation was assayed by phosphoimager analysis (lower panel) and the material loaded in the gel controlled by Comassie blue staining (top panel). **B:** Purified CotB (2 μM) incubated with purified CotH or CotH$^{D228Q}$ (0.5 μM) in the presence of [γ-32P]ATP and the reaction was stopped with loading buffer at the indicated times. CotH or CotB alone were used as control. In the panels one right, staurosporine (STP) was added at the indicated concentration to the phosphorylation reaction. The reactions were run in a SDS-PAGE, phosphorylation was assayed by phosphoimager analysis (lower panel) and the material loaded in the gel controlled by Coomassie blue staining (top panel). The position of molecular weight markers (in kDa) is shown on the left side of the panels.

**Figure S6 – Site of CotB and CotG phosporylation.** Whole cell extracts were prepared from a wild type strain, a *cotB* (B-) or a *cotG* (G-) mutant, at the indicated time (in hours) after the initiation of sporulation in DSM. The extracts were fractionated by centrifugation into a forespore and a mother cell fraction, and processed for immunoblotting with the indicated antibodies. The samples from the B- and G- strains were only analysed at hour 10 of sporulation. Arrows show the position of CotB-66 (blue), CotB-46 (red), and CotG (black); the asterisk represent likely multimeric forms of CotB, and the parenthesis indicates likely degradation products of $\sigma^A$. Molecular weight markers (M) in kDa, are shown on the left side of the panels.

**Figure S7 – Construction and analysis of a *cotB*$^{\Delta SKR}$ mutant. A:** A WT copy of *cotB* or an allele bearing an in-frame deletion of the $SKR^B$-coding region (*cotBΔSKR*), was introduced at the *amyE* locus of a *cotB* in-frame deletion mutant. The extent of the deletion is shown by the broken dashed line. **B**: diagram showing the expected molecular weight for the forms of CotB produced from the WT and *cotBΔSKR* alleles, and also of a form, CotB$^{P}$ that may arise through proteolytical cleavage before the start of the SKR$^{B}$ region. **C:** Spores were purified from DSM cultures of the *ΔcotB* mutant bearing either the WT allele or *cotBΔSKR* alleles at *amyE*. The coat proteins were extracted using an SDS/DTT regime, electrophoretically resolved and the gel stained with Coomassie (top panel) or subject to immunoblotting with an anti-CotB antibody (bottom). CotB-66, blue arrow; CotB-46, red arrow, CotB$^{SKR}$, green arrow. Asterisks indicate possible degradation products. The position of molecular weight markers (in kDa) is shown on the left side of the panels. A band at about 46 kDa seen by immunoblotting in the *cotBΔSKR* strain (black arrow in the bottom panel) may be a cross-reactive species that is more extractable in this mutant.

### Supplemental Tables

#### Table S1. Bacterial strains.

| Strain | Relevant Genotype/Phenotype[a] | Origin/ Reference |
|---|---|---|
| ***E. coli*** | | |
| BL21(DE3) | *fhuA2 [lon] ompT gal (λ DE3) [dcm] ΔhsdS λ DE3 = λ sBamHIo ΔEcoRI-B int::(lacI::PlacUV5::T7 gene1) i21 Δnin5* | Lab stock |
| DH5α | *ΔlacZ Δ M15 Δ(lacZYA-argF) U169 recA1 endA1 hsdR17(rK-mK+) supE44 thi-1 gyrA96 relA1* | “ |
| AH4051 | BL21(DE3) pMS16 (CotB::His$_6$) / Km$^R$ | “ |
| AH4071 | AH4051 pAP39 (CotG) / Cm$^R$ Km$^R$ | This work |
| AH4072 | BL21(DE3) pAP39 (CotG) / Cm$^R$ | “ |
| AH4083 | AH4051 pNP5 (CotH$^{D228Q}$-*strep-tag*) / Cm$^R$ Km$^R$ | “ |
| AH4084 | AH4051 pNP6 (CotG+CotH$^{D228Q}$-*strep-tag*) / Cm$^R$ Km$^R$ | “ |
| AH4085 | BL21(DE3) pNP5 (CotH$^{D228Q}$-*strep-tag*) / Cm$^R$ | “ |
| AH4086 | BL21(DE3) pNP6 (CotG+CotH$^{D228Q}$-*strep-tag*) / Cm$^R$ | “ |
| AH4102 | BL21 (DE3) pCD10 (CotH$^{WT}$-*strep-tag*) / Cm$^R$ | “ |
| AH4104 | AH4051 pCD10 (CotH$^{WT}$-*strep-tag*) / Cm$^R$ Km$^R$ | “ |
| ***B. subtilis*** | | |
| MB24 | *trp*C2 *met*C3/wild-type, Spo$^+$ | Lab stock |
| PY79 | Prototrophic derivative of *B. subtilis* subsp. *subtilis* 168 | “ |
| AH141 | *trpC2 metC3 ΔcotB::cat*/ Cm$^R$ | “ |
| AH1103 | *trpC2 metC3 ΔcotH::cat*/ Cm$^R$ | “ |
| AH1497[b] | trpC2 metC3 *cotG*ΩpMS43/Cm$^R$ | “ |
| AH4097 | trpC2 *metC3 ΔcotH::cat ΔamyE::cotH$^{wt}$* / Cm$^R$ Nm$^R$ | This work |
| AH4098 | *trpC2 metC3 ΔcotH::cat ΔamyE::cotH$^{D228Q}$* / Cm$^R$ Nm$^R$ | “ |
| AH2195 | *trpC2 metC3 ΔcotXYZ* | Lab stock |
| AH6992 | *trpC2 metC3 ΔcotG* | This work |
| AH7002 | *trpC2 metC3 ΔcotG ΔcotXYZ*/ Nm$^R$ | “ |
| AH7001 | *trpC2 metC3 ΔcotB ΔcotG* | “ |
| AH6988 | PY79 with *cotZ-gfp* | Lab stock |
| AH6990 | *ΔcotG cotZ-gfp* | This work |
| AH6899 | *ΔcotB cotZ-gfp* | “ |
| AH7000 | *ΔcotH cotZ-gfp* | “ |
| AH6986 | *cotH$^{WT}$ cotZ-gfp* | “ |
| AH6987 | *cotH$^{D228QT}$ cotZ-gfp* | “ |
| AH6997 | *trpC2 metC3 ΔcotB cotB$^{WT}$* | “ |
| AH6998 | *trpC2 metC3 ΔcotB cotB$^{SKR}$* | “ |

[a] Km, kanamycin; Cm, cloramphenicol; Nm, neomycin.

[b] The omega symbol (Ω) denotes that the plasmid was integrated into the *B. subtilis* chromosome by a single crossover event in the region of homology (a Campbell-type integration event).

#### Table S2. Oligonucleotides used in this study.

| Primer | Sequence (5´to 3´)[a] |
|---|---|
| cotB/5/Nco | GAGCCATGGGAATGAGCAAGAGGAGA |
| cotB/OMO/3 | GCCTAGGATCCGGGCATCACTTTATC |
| cotB/OMO/6 | GCCTAGGATCCGATGCGAAGCACCTC |
| cotB/Nco/ID | GTAGATAATGCCATGGGCCATTATAC |
| cotG/5/Nco | GAGCCATGGAATTGGGCCACTATTCC |
| cotG/Bam/3 | TACCTCCGCCGGGATCCTATTGAAAC |
| cotG-R/NotI | GCGGCCGCTTATTTGTATTTCTTTTTGAC |
| cotH/5/Nde | CTAAGGAGGACATATGATGAAGAATC |
| cot3/H/Bam | CCGTGGATCCACCCATTTTCACGCAT |
| cotH/5/Bam | GTAAGGAGGGATCCGGATGAAGAAT |
| cotH/3/Xho | CGCGAATACCCTCGAGCACGCATTCA |
| *cotH*-R/*XhoI* | CTCGAGTTATTTTTCGAACTGCGGGTGGCTCCAAGCGCTTAAAATACTTAAATGA TCTTTGAGGTATTG |
| *cotH*-F/*NdeI* | CCGTTATCGCATATGAAGAACCAATCCAATTTACCGCTTTATCAGCTG |
| cotH95D | TTGAAAGCTTATGTACATAGCAACAACCGCC |
| cotH2140R | CCTAATTGTCCAGTTTTTCCGCGAATACCC |
| cotGcotHD | GCGAGATTTTTTGTGAGTGCGGTGCGAC |
| cotGcotHR | CACCGCACTCACAAAAAATCTCGCCGCAGCTATAAGAAGTCATGCCGC |
| cotH D228Q_D | CACCTCGAACTATCAAGGGTTTGTCCACAAC |
| cotH D228Q_R | GTTGTGGACAAACCCTTGATAGTTCGAGGTG |

[a] Restriction sites are underlined; base substitutions are shown in bold.

#### Table S3. Plasmids.

| Plasmid | Relevant features | Origin/reference |
|---|---|---|
| pACYCDuet-1 | T7*lac* expression vector | Novagen |
| pET-30a | T7*lac* expression vector | “ |
| pMS16 | *cotB* in pET30a(+)/Km | (Zilhao *et al.*, 2004) |
| pAP39 | *cotG* in pACYCDuet-1 | This work |
| pCD6 | Derivative of pMLK83 | “ |
| pCD7 | *cotH* in pCD6 | “ |
| pCD8 | *cotH*$^{D228Q}$ in pCD6 | “ |
| pCD10 | *cotH* in pACYCDuet-1 | “ |
| pCD11 | *cotH* in pAP39 | “ |
| pNP5 | *cotH*$^{D228Q}$ in pACYCDuet-1 | “ |
| pNP6 | *cotH*$^{D228Q}$ in pAP39 | “ |

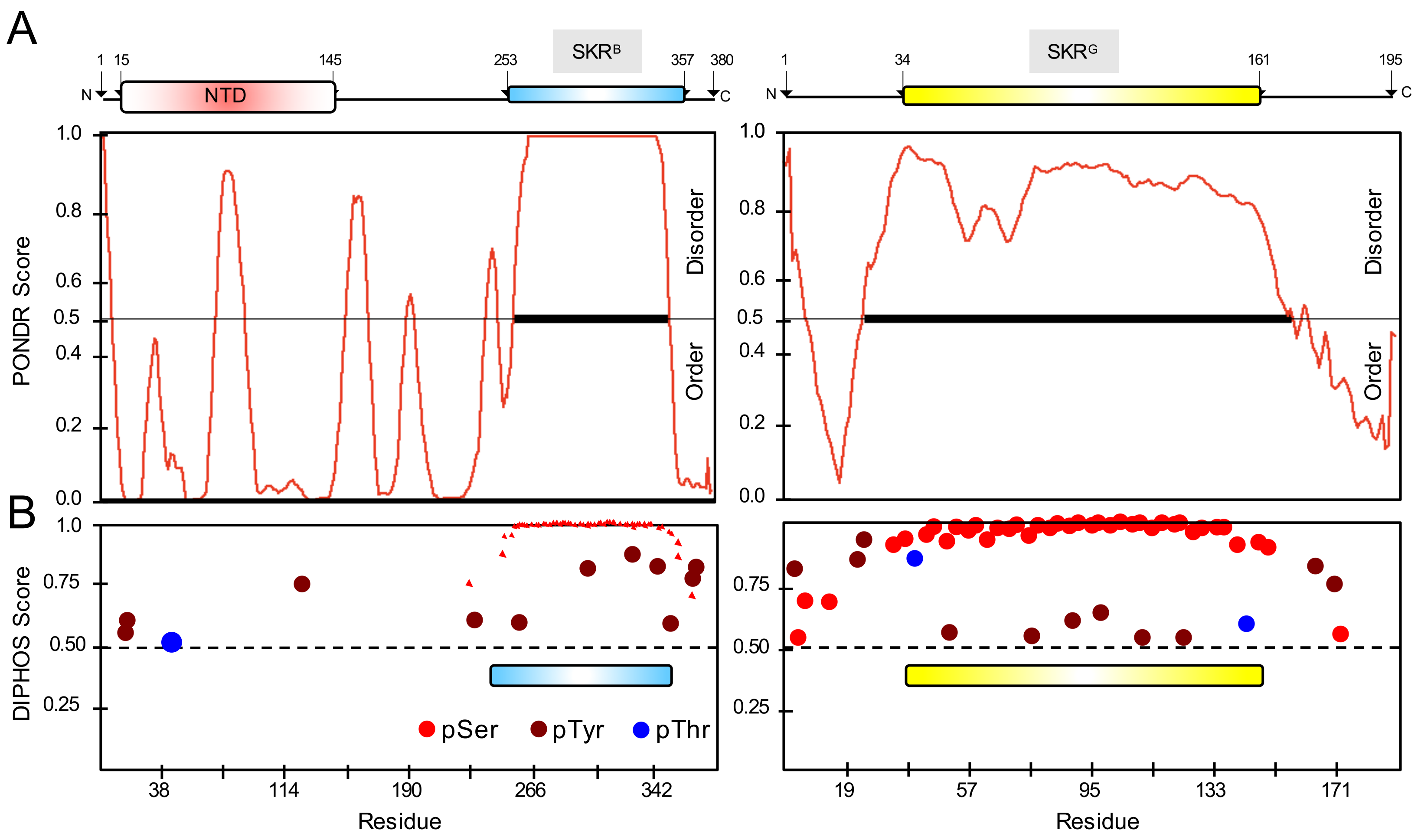
A
SKRB
SKRG
NTD
N
C
1
15
145
253
357
380
34
161
195
PONDR Score
1.0
0.8
0.6
0.5
0.4
0.2
0.0
Disorder
Order
B
DIPHOS Score
0.75
0.50
0.25
pSer
pTyr
pThr
38
114
190
266
342
19
57
95
133
171
Residue

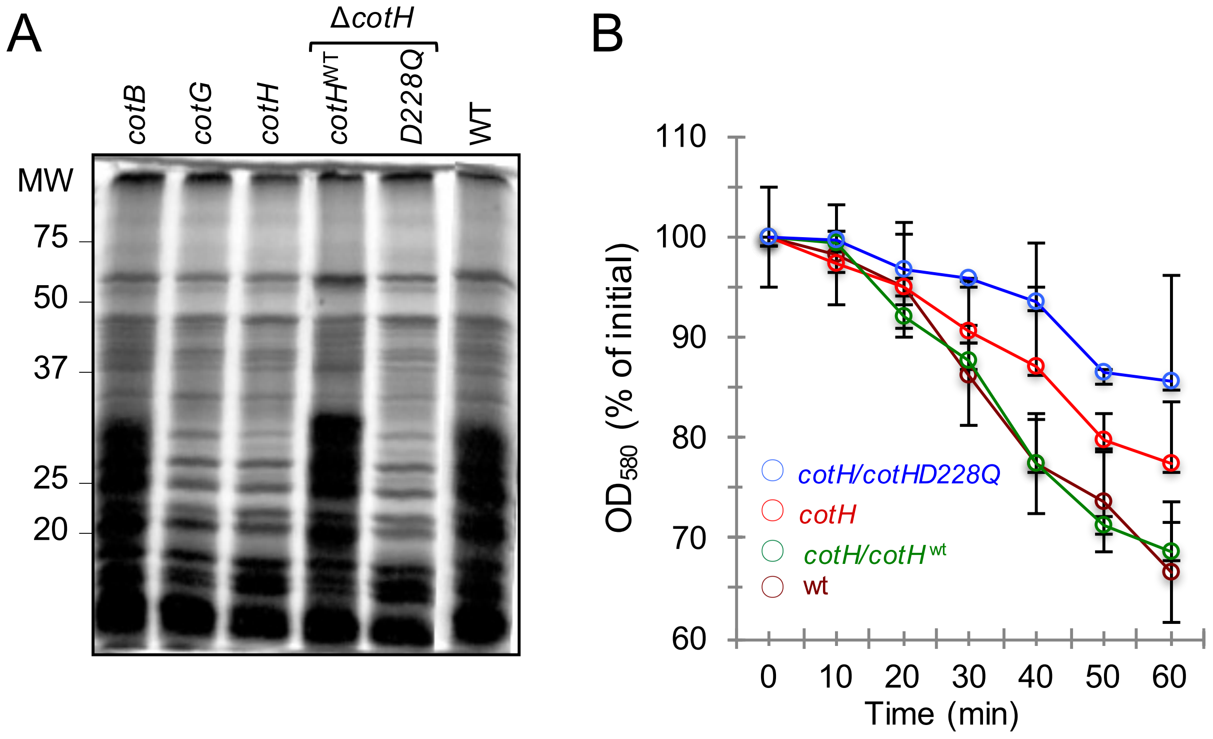
A
ΔcotH
cotB
cotG
cotH
cotHWT
D228Q
WT
MW
75
50
37
25
20
B
OD580 (% of initial)
110
100
90
80
70
60
0
10
20
30
40
50
60
Time (min)
cotH/cotHD228Q
cotH
cotH/cotHwt
wt

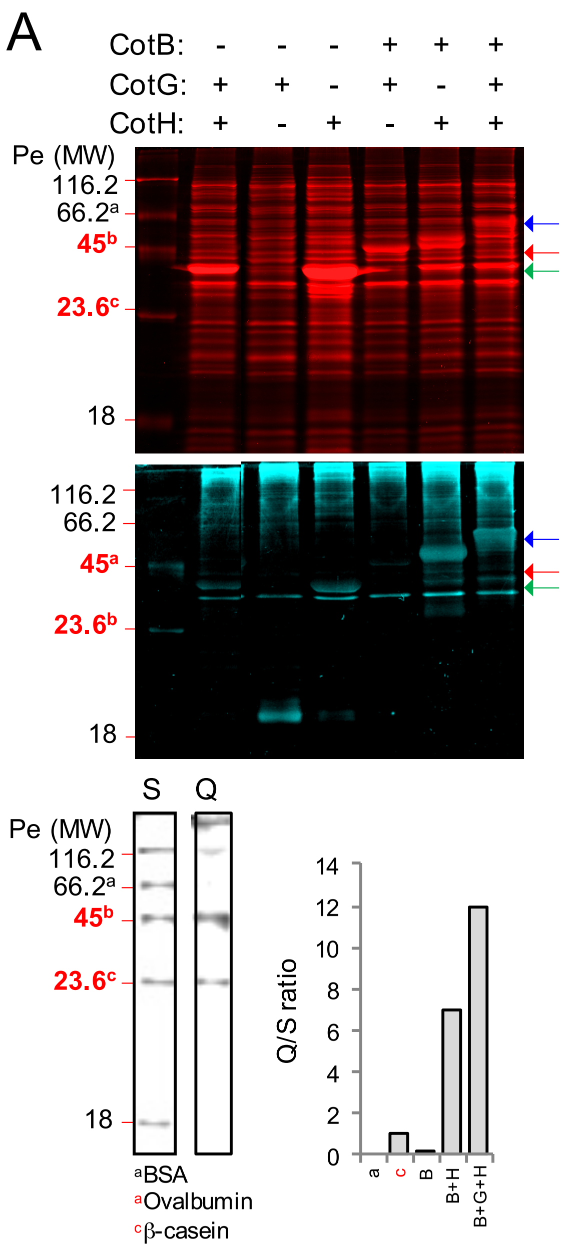
A
CotB: - - - + + +
CotG: + + - + - +
CotH: + - + - + +
Pe (MW)
116.2
66.2a
45b
23.6c
18
116.2
66.2
45a
23.6b
18
S
Q
Pe (MW)
116.2
66.2a
45b
23.6c
18
aBSA
aOvalbumin
cβ-casein
Q/S ratio
14
12
10
8
6
4
2
0
a
c
B
B+H
B+G+H

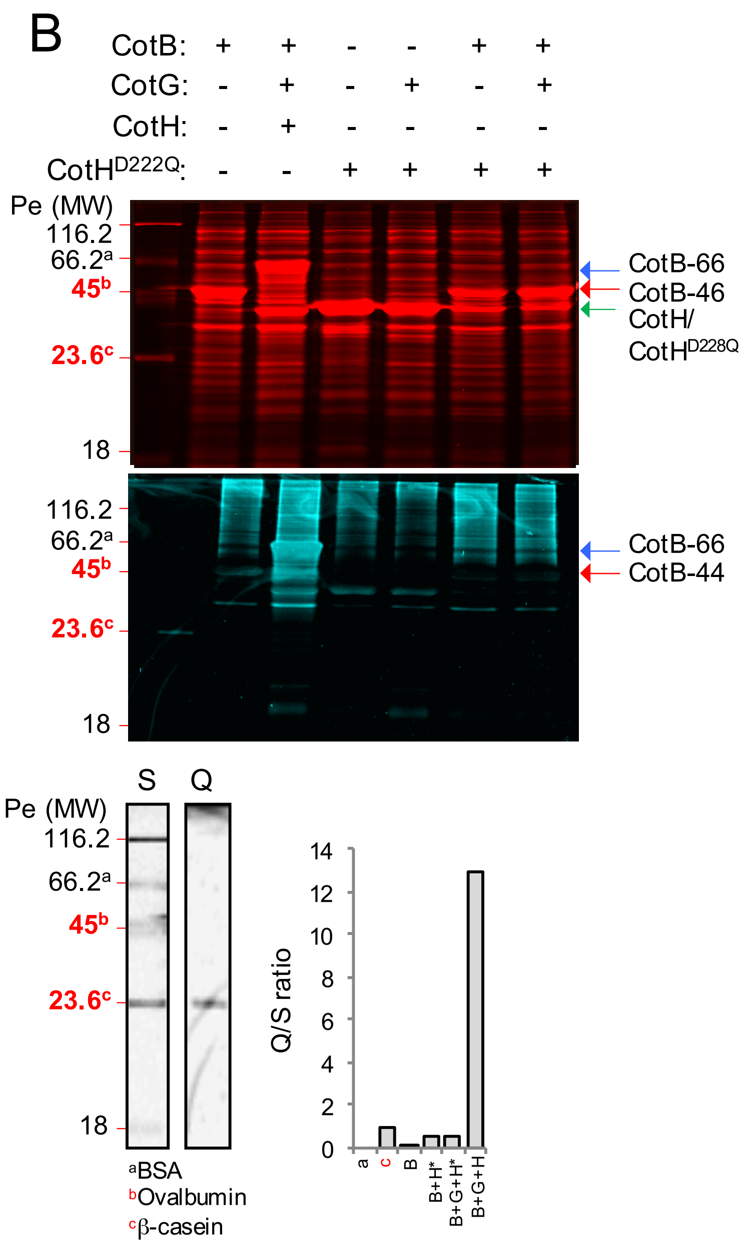
B
CotB: + + - - + +
CotG: - + - + - +
CotH: - + - - - -
CotHD222Q: - - + + + +
Pe (MW)
116.2
66.2a
45b
23.6c
18
CotB-66
CotB-46
CotH/
CotHD228Q
116.2
66.2a
45b
23.6c
18
CotB-66
CotB-44
S
Q
Pe (MW)
116.2
66.2a
45b
23.6c
18
aBSA
bOvalbumin
cβ-casein
Q/S ratio
14
12
10
8
6
4
2
0
a
c
B
B+H*
B+G+H*
B+G+H

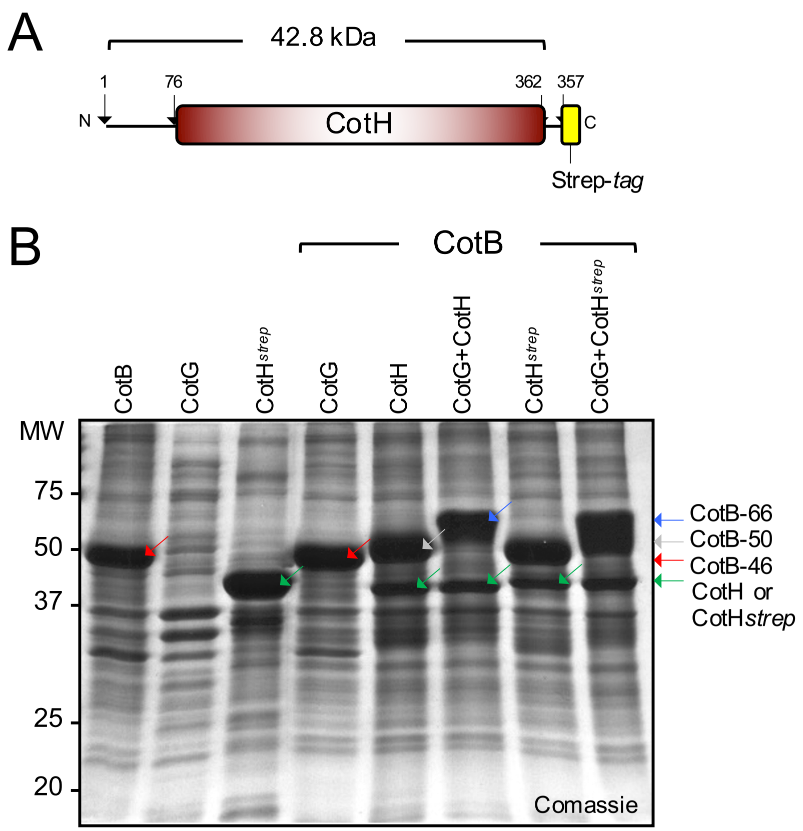
A
42.8 kDa
1
76
362
357
N
CotH
C
Strep-tag
B
CotB
CotB
CotG
CotHstrep
CotG
CotH
CotG+CotH
CotHstrep
CotG+CotHstrep
MW
75
50
37
25
20
Comassie
CotB-66
CotB-50
CotB-46
CotH or
CotHstrep

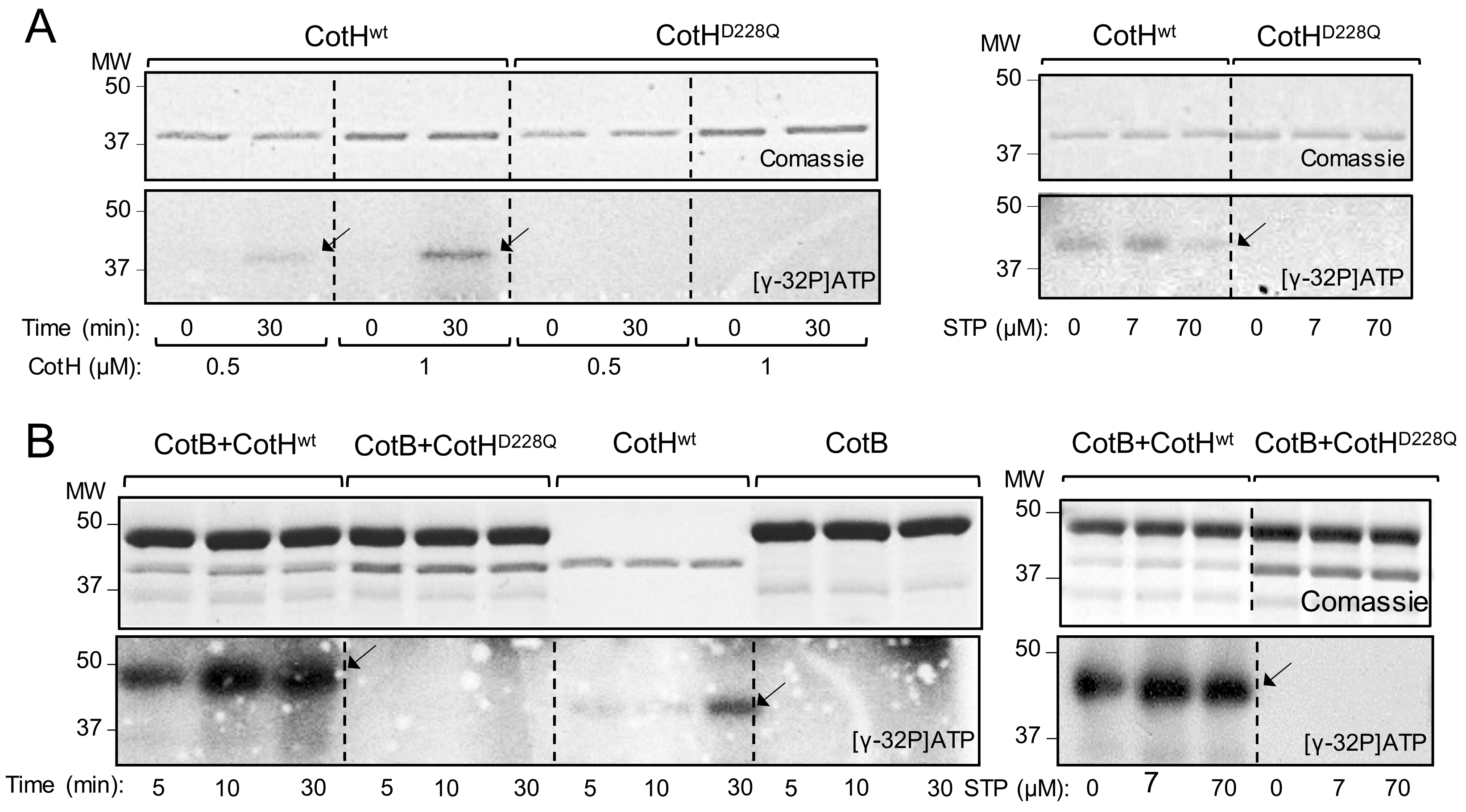
A
CotH^wt
CotH^D228Q
MW
50
37
Comassie
[γ-32P]ATP
Time (min): 0 30 0 30 0 30 0 30
CotH (μM): 0.5 1 0.5 1
STP (μM): 0 7 70 0 7 70
B
CotB+CotH^wt
CotB+CotH^D228Q
CotH^wt
CotB
Time (min): 5 10 30 5 10 30 5 10 30 5 10 30
STP (μM): 0 7 70 0 7 70

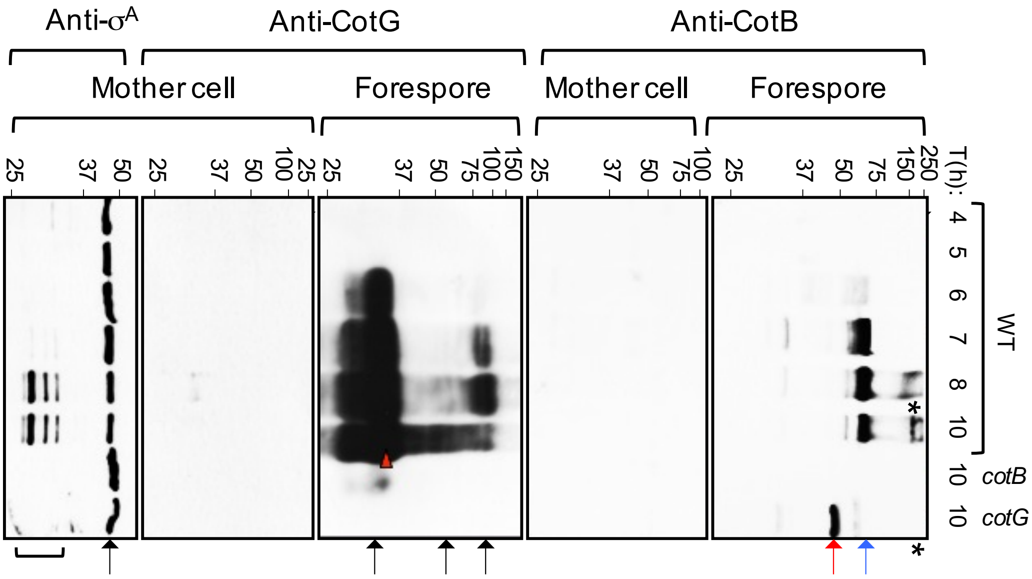


Freitas *et al.* - Fig. S6

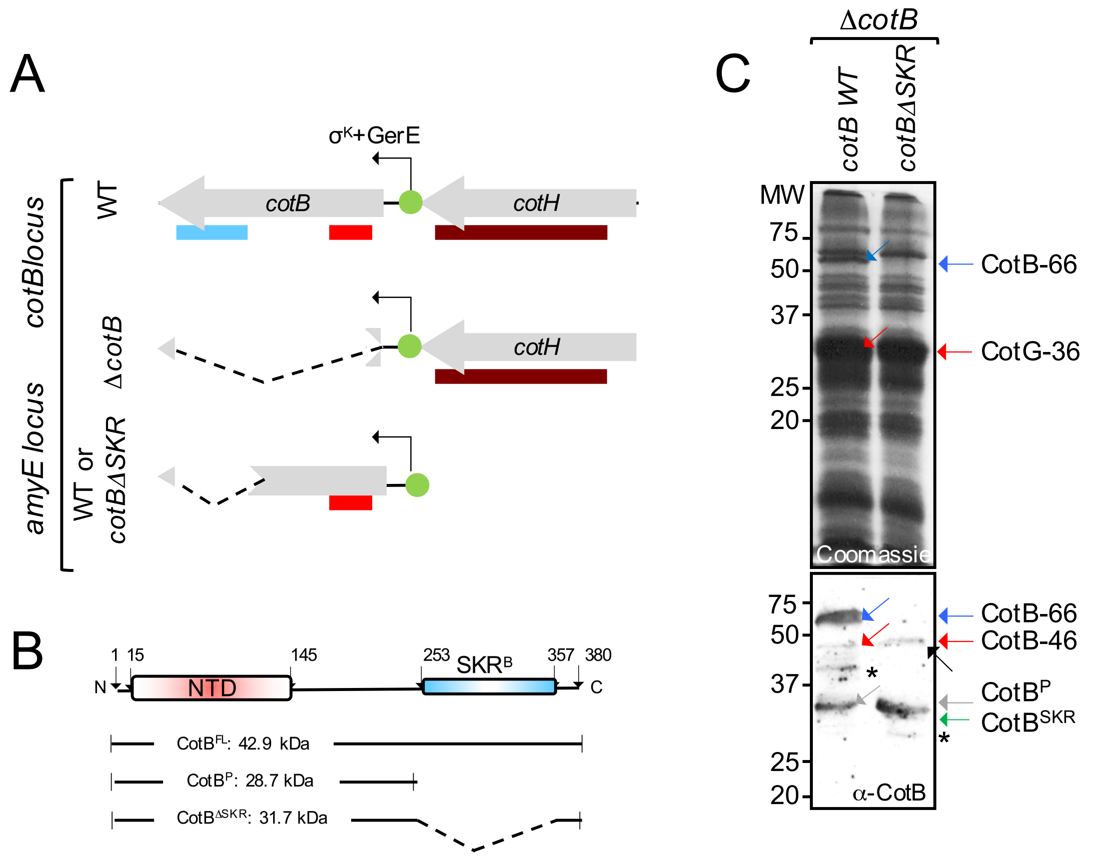
A
cotBlocus
amyE locus
WT
ΔcotB
WT or
cotBΔSKR
σK+GerE
cotB
cotH
cotH
B
N
1 15
NTD
145
253
SKRB
357 380
C
CotBFL: 42.9 kDa
CotBP: 28.7 kDa
CotBΔSKR: 31.7 kDa
C
ΔcotB
cotB WT
cotBΔSKR
MW
75
50
37
25
20
Coomassie
CotB-66
CotG-36
75
50
37
25
20
α-CotB
CotB-66
CotB-46
CotBP
CotBSKR
